# Trajectories from Snapshots: Integrated proteomic and metabolic single-cell assays reveal multiple independent adaptive responses to drug tolerance in a BRAF-mutant melanoma cell line

**DOI:** 10.1101/767988

**Authors:** Yapeng Su, Guideng Li, Melissa E. Ko, Hanjun Cheng, Ronghui Zhu, Min Xue, Jessica Wang, Jihoon W. Lee, Luke Frankiw, Alexander Xu, Stephanie Wong, Lidia Robert, Kaitlyn Takata, Sui Huang, Antoni Ribas, Raphael Levine, Garry P. Nolan, Wei Wei, Sylvia K. Plevritis, David Baltimore, James R. Heath

## Abstract

The determination of individual cell trajectories through a high-dimensional cell-state space is an outstanding challenge, with relevance towards understanding biological changes ranging from cellular differentiation to epigenetic (adaptive) responses of diseased cells to drugging. We report on a combined experimental and theoretic method for determining the trajectories that specific highly plastic BRAF^V600E^ mutant patient-derived melanoma cancer cells take between drug-naïve and drug-tolerant states. Recent studies have implicated non-genetic, fast-acting resistance mechanisms are activated in these cells following BRAF inhibition. While single-cell highly multiplex omics tools can yield snapshots of the cell state space landscape sampled at any given time point, individual cell trajectories must be inferred from a kinetic series of snapshots, and that inference can be confounded by stochastic cell state switching. Using a microfludic-based single-cell integrated proteomic and metabolic assay, we assayed for a panel of signaling, phenotypic, and metabolic regulators at four time points during the first five days of drug treatment. Dimensional reduction of the resultant data set, coupled with information theoretic analysis, uncovered a complex cell state landscape and identified two distinct paths connecting drug-naïve and drug-tolerant states. Cells are shown to exclusively traverse one of the two pathways depending on the level of the lineage restricted transcription factor MITF in the drug-naïve cells. The two trajectories are associated with distinct signaling and metabolic susceptibilities, and are independently druggable. Our results update the paradigm of adaptive resistance development in an isogenic cell population and offer insight into the design of more effective combination therapies.

## Introduction

Cellular processes ranging from the development of drug-tolerant states in cancer cells to stem cell differentiation, can be described as cell state changes. Specifically, certain cancer cells that are initially responsive to targeted inhibitors that act against these oncogenic drivers^1–4^ can evolve into a drug-tolerant state via non-genetic mechanisms, perhaps preceding the emergence of drug-resistant clones^5–7^. The molecular details of how the cancer cells transition between the two states can inform the use of additional drugs designed to arrest the transition^8–10^. Dating back to the epigenetic landscapes of Waddington^11^, a prevalent picture is that cells take a single path that connects the initial to the final state, but this does not have to be the case. In fact, if cells can take multiple independent paths between the two states, then the challenge of finding drug combination that can arrest the unfavorable cell state transition is significantly increased. Here we investigate a highly plastic cancer cell line that, when treated with a targeted inhibitor, switches from a rapidly dividing, drug responsive state to a drug-tolerant, slow cycling state within a few days. We show that the cells can take multiple classes of trajectories between the two states. Each trajectory class is characterized by a unique signaling and metabolic networks with distinct druggable susceptibilities. From a functional perspective, cell state changes are often accompanied by changes in gene expression^10,12–15^, protein signaling^12,14,16–22^ and cellular metabolism.^23–26^ Highly multiplex single-cell methods^27–30^ can provide powerful tools for mapping out cell-state landscapes associated with cell state changes^20,31–35^. However, capturing the trajectories that individual cells take as they traverse those landscapes is challenging, even for the case of an isogenic cell line. This is because multiplex single-cell omics methods only provide snapshots of the occupied cell state space at a given instant. Measured similarities between cells captured at successive time points can imply probable paths through the landscape^36–40^. However, cells may stochastically switch from one state to another, so an individual cell may not take a smooth trajectory between states. Time-lapse imaging methods can map individual cell trajectories, but for only a couple of analytes for each cell, and so provide a limited view of the cell state space^41–43^. Thus, the ability to extract cellular trajectories from a kinetic series of cell state space snapshots would have high value. Here we report on combined experimental and theoretical approaches towards addressing this fundamental challenge.

We utilized a patient-derived *BRAF^V600E^* mutant melanoma cancer cell line as a model for the rapid development of drug tolerance against targeted inhibitors. Under BRAF inhibition, these highly plastic cells rapidly (and reversibly) transition from a drug-responsive state to a drug-tolerant state^12,19^. We characterized this transition using integrated single-cell functional proteomic and metabolic assays designed to broadly sample proteins and metabolites associated with selected cancer hallmarks and cell state-specific processes. Dimensional reduction, information-theoretic analysis, and visualization of the time-series single-cell data uncovered a complex cell state space landscape, and hinted at the possibility of two distinct paths between drug-naïve and drug tolerant states. Further experiments tested whether these paths constituted independent (and thus independently druggable) cellular trajectories. In fact, we find that even isogenic tumor cells can undertake different, independent trajectories to drug tolerance. The two trajectories are associated with distinct signaling and metabolic networks, and are independently druggable. This finding challenges the current paradigm of targeted inhibitor resistance development, and also provides guidelines for assessing the value of combination therapies.

## Results

### Integrated single-cell proteomic and metabolic analysis characterizes early BRAFi adaptation in melanoma cells

We characterized drug adaptation in individual melanoma cells by assaying for a panel of selected proteins, plus glucose uptake, in BRAF*^V600E^* mutant M397 cell cultures during the first five days of BRAFi treatment using the Single Cell Barcode Chip (SCBC)^12,20,29,44–47^ (Fig 1a). Following 0, 1, 3 and 5 days (D0 control, D1, D3, and D5) of drug treatment, individual cells were isolated into nanoliter-volume microchambers within an SCBC. Each isolated cell was lysed *in situ* to release the cellular contents. Each microchamber within an SCBC contains a full barcode array in which each barcode element is either an antibody for specific protein capture^48^ or a molecular probe designed to assay for a specific metabolite via a competition assay^46,47^ (Fig.1a). The design of this panel was informed by transcriptomic analysis of BRAFi-treated M397 cells (Supplementary Fig. 1) and existing literature^12,14,16,23,49,50^. The panel broadly samples various functional and metabolic hallmarks of cancer, as well as cell state markers.

**Fig. 1.**
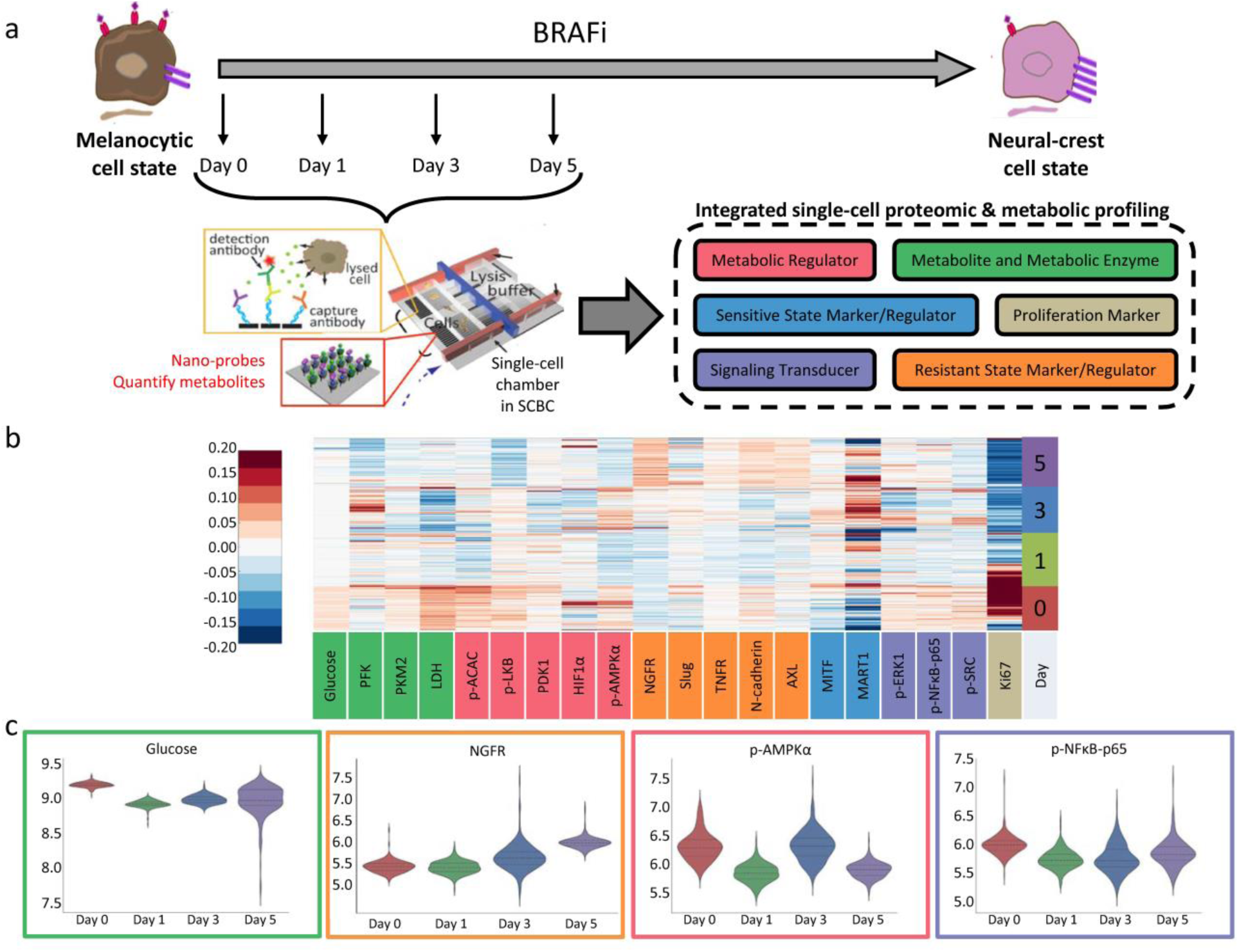
Single-cell proteomic and metabolic analysis of early drug response in M397 cells. **a.** The single-cell integrated proteomic and metabolic analysis experiments design. Cells from different time points during BRAFi treatment are harvested and individually analyzed using the microfluidic based single-cell barcode (SCBC) technology. Each cell was characterized for the levels of 6 different categories of markers. **b.** Heatmap representation of integrated proteomic and metabolic analysis dataset. Each row represents an individual cell and each column (except the last column) represents an individual analyte, with the color in the heatmap representing measured level of the analyte. The last column represents the number of days after starting BRAFi treatment. On the X-axis, markers are colored corresponding to which of the six functional categories they belong to. **c.** Violin plot representation of distribution of certain representative markers across 4 time points. Y-axis represents natural log of measured marker level. Each plot is bordered by the color of the functional category of the measured marker.

Single-cell profiling of BRAFi-naïve (D0) M397 cells revealed heterogeneous levels of many assayed markers at baseline. Referring to Fig. 1b,c **and** Supplementary Fig. 2, certain analytes exhibit high variability across the cell population. These include the melanocytic lineage transcription factor MITF and its downstream melanocytic cell state marker MART1, the metabolic regulators HIF1α and p-AMPKα, and the proliferation marker Ki67. The variance in Ki67 implies that the population contains both rapid-cycling and slow-cycling cells. By contrast, a high glucose uptake and the expression of metabolic enzymes LDH and PKM2 were relatively uniform from cell-to-cell. Drug treatment initially (at D1) inhibits glucose uptake and represses most metabolic regulators and signaling phosphoproteins, as well as Ki67. The repression of these cancer hallmarks reflects blockage of the key oncogenic signaling pathway upon initial BRAF inhibition. The drug also promotes transient cell differentiation followed by dedifferentiation, as evidenced by an increase of MART1 expression in D3 followed by its downregulation in D5. However, a small subpopulation of M397 cells remains Ki67-high in D1, implying a slower drug response in that subset of cells. At D3, most analytes exhibit a sharp and transitory increase in variance, which shrinks by D5. This change includes all of the metabolic regulators except p-LKB, all resistant state markers and regulators except Slug, all of the metabolic enzymes, and all of the signaling phosphoproteins. The increased magnitude of the fluctuations of many markers at D3, based upon previous reports^45,51^, implies one or more cell state changes near this time point. By D5, glucose uptake has increased back to near D0 levels, but with increased variance. Ki67 is further decreased, and with sharply decreased variance relative to D0. Additionally at this day, the variance and abundance of the epithelial–mesenchymal transition (EMT)-related transcription factor, Slug, has increased, indicating the emergence of some cells that are trending towards a mesenchymal phenotype. Further, the levels of the other assayed protein markers that are associated with drug resistance (AXL, N-cadherin, NGFR, and TNFR) are all higher by D5. The upregulation of glucose uptake and many resistance marker indicates that cells have initiated drug resistance programs by D5. Thus, single-cell integrated proteomic and metabolic analysis, when viewed at the level of individual analytes, provides evidence of initial drug response at D1, a drug-induced cell state change at D3, and emerging drug tolerance at D5, prior to an increase in cell proliferation (full drug resistance) which has been shown to occur a few weeks later. These observations are all consistent with existing literature^12,14,16,52,53^.

### Dimensional reduction analysis implies multiple trajectories towards drug adaptation

Simultaneous visualization of the time-dependent, coordinated changes across multiple markers requires algorithms that can reduce the high-dimensionality of the dataset. We applied two such algorithms: the FLOW-MAP algorithm^54^ and the t-SNE algorithm^32^. Both approaches provided an intuitive representation of the dataset (Fig. 2 **and** Supplementary Figs. 3 and 4). FLOW-MAP analysis revealed that melanoma cells clustered primarily based upon drug exposure time (Fig. 2**, upper left plot**) indicating chronological cell state trajectories. Most untreated M397 cells (in the lower left of the graph) were characterized by uniform levels of all measured analytes excepting N-cadherin, MITF, HIF1α, Ki67 and MART1 (see analyte-specific plots of Fig. 2 **and** Supplementary Fig. 3). Most of these non-uniformly expressed proteins exhibit differences that vary gradually from left-to-right across the D0 cluster of cells, with a small subpopulation of untreated cells (right hand side of D0 cluster) exhibiting lower expression of Ki67, MITF, and MART1. These features point to a small group of dedifferentiated, slow-cycling cells. Upon BRAFi treatment, the cells initially split to occupy two regions of the FLOW-MAP. At D1 (green points), the majority of the cells cluster to the upper right of the D0 cells, while a small subpopulation clusters directly to the right of the D0 group. This trend continues at D3, with most cells clustering above the largest D1 mass, while a small number cluster to the right of the small D1 group. By D5 (purple), all cells cluster to the right hand side of the graph. The bifurcation of cells at Day 1 and 3 implies the possibility of “upper” and “lower” trajectories towards the drug-tolerant state. The possibility of two classes of trajectories was also indicated by t-SNE analysis^32^ (Supplementary Fig.4). Thus, both computational analyses of the single-cell data set indicate a bifurcated drug response during the early stages of BRAFi adaptation.

**Figure. 2.**
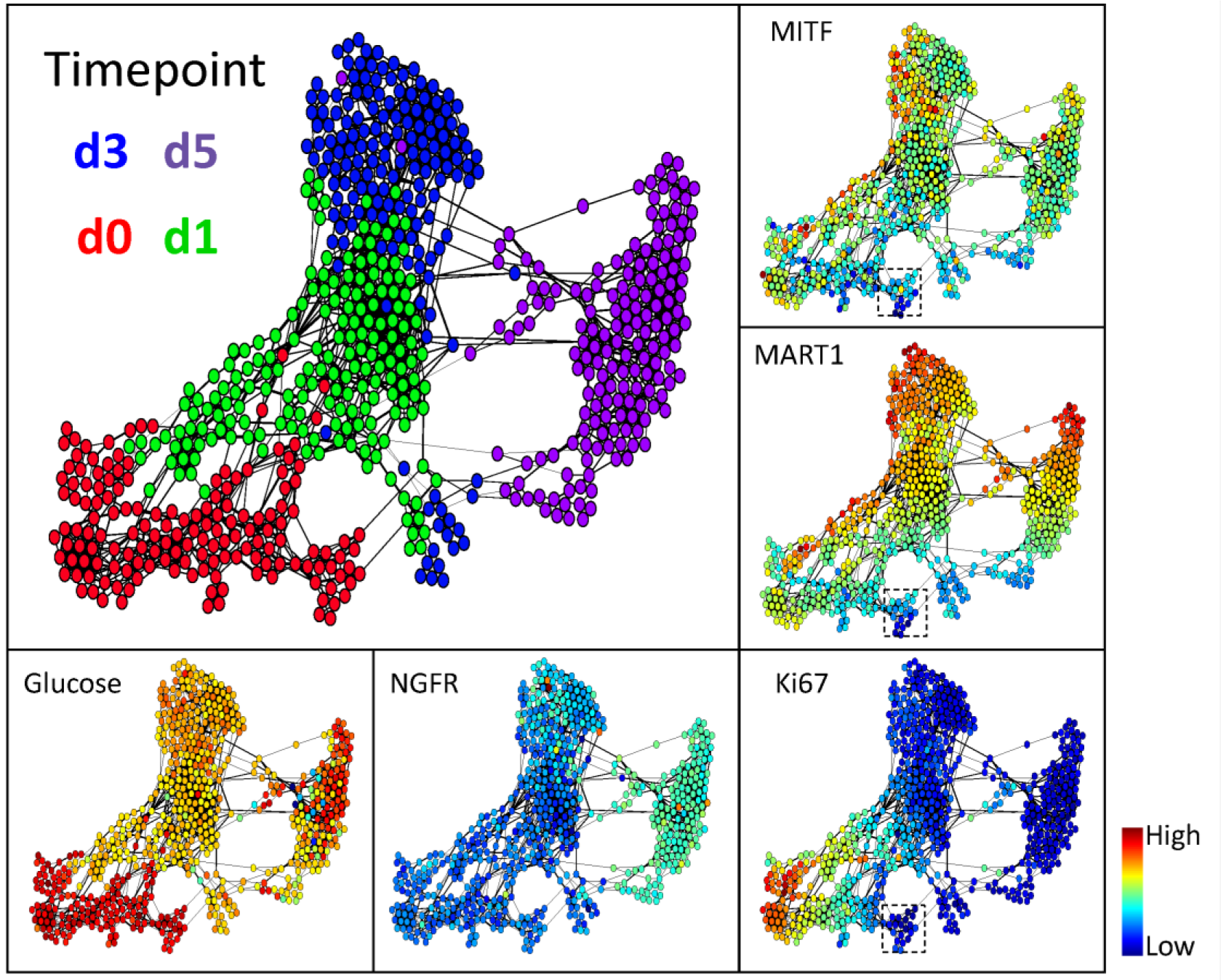
Visualization of single-cell data by FLOW-MAP. Each dot represents an individual cell. The distance between each pair of cells represents the overall multi-omic dissimilarity between them. Cell pairs that are close enough are linked with an edge in between. The colors of the dots in the main panel (upper-left) represent BRAFi exposure time (0, 1, 3, or 5 days) of the corresponding cells. Dot colors in the other panels represent the abundance of each marker in each cell. The dashed-line box in the panels for MITF, MART1, and Ki67 levels shows a small subpopulation of day-0 cells that are slow cycling with less melanocytic phenotype.

### Surprisal analysis uncovers analyte modules of the bifurcated drug-response trajectories

To further dissect the dynamics of molecular changes associated with the bifurcated drug-response trajectories, we applied surprisal analysis^55–57^ to our single-cell dataset. Surprisal analysis is a thermodynamics-inspired method that has been broadly applied to understanding large-scale bulk and single-cell omics data sets^51,55,57–59^. This approach is based on the identification of the steady state of the system (formally speaking the state of minimum free energy), and any constraints (analyte modules) that increase the free energy from this theoretical minimum^57,60^. Using this approach, we identified two main modules, each representing a set of analytes that are coordinately changing together across cells. The predicted expression of all 20 analytes based on these two modules matched well with the measured single-cell dataset (Supplementary Figs. 5 and 6), demonstrating that modules 1 and 2 recapitulate the overall changes of all molecular signatures across all cells over the five-day course of drug treatment.

The influence score (the lambda values defined in ref ^57^) of a module in a cell represents the extent to which the module-associated analytes are enriched or repressed in that cell. Modules 1 and 2 were visualized by color-coding their influence scores onto each node in the FLOW-MAP graph (Fig. 3a). We found that the influence score of module 1 gradually increased from negative (blue) to positive (red) value along both the upper and lower paths, with a clear “biophysical barrier” (lambda1 = 0) in the middle time points (Fig. 3a, **left panel**). We have previously shown that such a sign change can imply a cell state transition^55^. The time-dependence of module 1 appears to reflect the transition from a drug-responsive state to a slow-cycling, drug-tolerant state between days 1 and 3. This observation is consistent with the negative correlation of Ki67 expression and positive correlation of NGFR/AXL expression with the module 1 score (Supplementary Fig. 7). The module 2 projection on the FLOW-MAP also exhibits a sign change, or biophysical barrier (lambda2 = 0), which separates the upper and lower paths (Fig. 3a**, right panel**). In fact, module 2 distinguishes cell subpopulations for each of the analyzed time points. Notably, the expression of melanocytic phenotype transcription factor MITF and its downstream protein MART1 both showed negative correlations with module 2 score (Fig 3b **and** Supplementary Fig. 8), indicating that the separation of the two paths may be related to the melanocytic lineage of the cells. In summary, surprisal analysis resolves both time-dependent and path-specific modules. It also reveals that, as the cells advance from drug-naïve to drug-tolerant, they occupy a rather complex landscape of cellular states separated by multiple biophysical barriers (Supplementary Fig. 9).

**Fig. 3.**
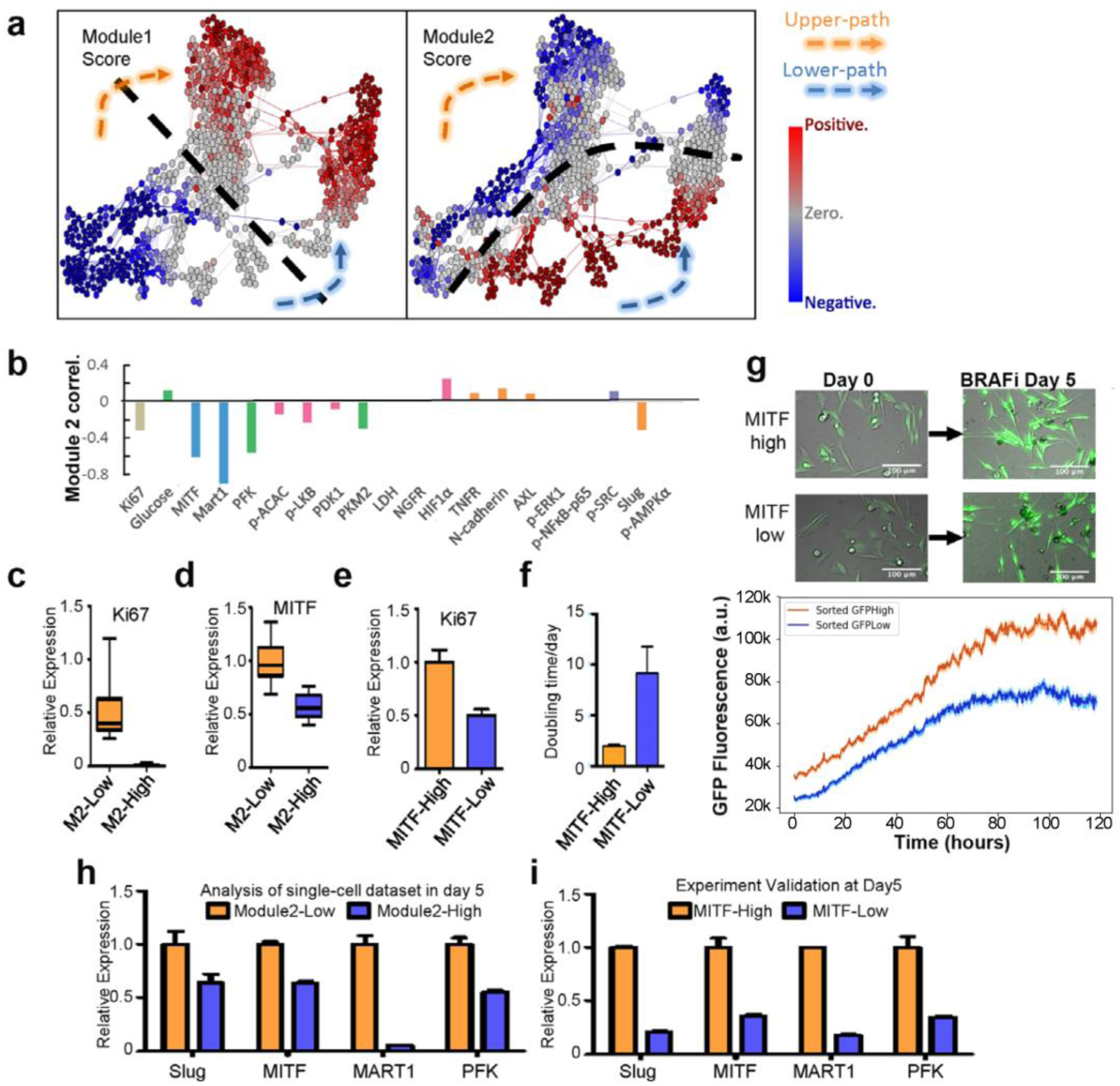
Surprisal analysis identifies time-dependent and path-specific analyte modules that explain the bifurcated trajectories and identify MITF as a transcription factor regulating the bifurcation. **a.** Visualization of the influence score of the two regulatory modules identified from surprisal analysis. Module1 is time-dependent, while module2 exhibits a path-specific pattern. The dashed black lines indicate the region for which the respective module scores of each cell approach zero. **b.** Pearson correlation between individual marker levels and the module2 score. **c, d.** Ki67 and MITF expression level in module2 score-high and -low subpopulations at day 0. **e.** Ki67 relative expression, measured by q-PCR in sorted MITF-high and MITF-low cells at day 0. **f.** Doubling time measured in treatment-naïve condition, collected from sorted MITF-high and MITF-low cells at day 0. **g.** Single-cell time-lapsed microscopy analysis of MITF-activity during 5 days of BRAFi. Top panel: Time-lapse images of sorted GFP-High and GFP-low cells before and after 5 days of BRAFi. Bottom panel: Average trace representing MITF activity dynamics across single MITF-Low (blue trace) and MITF-High (orange trace) cells over 5 days of BRAFi. Shading indicates SEM of the mean. **h.** Slug, MITF, MART1 and PFK relative expression levels in module2 score-high and -low subpopulations, collected from cells at day 5 and analyzed from single-cell dataset. **i.** Slug, MITF, Mart1 and PFK expression, measured by q-PCR in sorted MITF-high and MITF-low day-0 cells that have been treated with BRAFi for 5 days.

### Experimental validation supports bifurcated drug-response trajectories

Surprisal analysis provides theoretical support for the existence of both the upper and lower paths from drug-naïve to drug-tolerant cell states. However, experimental validation is required to determine whether individual cells exclusively follow a single trajectory along one path or the other, or if cells stochastically switch between paths. The map of module2 on the D0 cells data hints at biological differences that separate even the untreated D0 cells into two subpopulations (Supplementary Fig. 9). The expression levels of the transcription factor MITF and its direct downstream target MART1 are among the top four markers that distinguish the two D0 subpopulations (Supplementary Fig. 10). This finding suggests that drug-treated MITF^low^ cells might follow the lower path, while MITF^high^ cells might follow the upper path (Supplementary Figs. 11 a). We thus generated MITF-GFP reporter cell lines and sorted GFP^high^ (MITF^high^) and GFP^low^ (MITF^low^) subpopulations (Supplementary Figs. 11 b and 12). Consistent with our hypothesis, MITF^high^ cells displayed higher expression of Ki67 and MITF as well as a shorter doubling time relative to sorted MITF^low^ subpopulations (Fig. 3 c-f). This data is consistent with reported observations of melanoma phenotype switching from a melanocytic, highly proliferative state to a non-melanocytic, more invasive state^61^. It also confirmed that the two subpopulations in D0 cells can be separated using this reporter system, and further suggests that the MITF^high^ and MITF^low^ subpopulations at D0 may represent cells destined to follow the upper and lower paths, respectively, following drug treatment.

To quantify the frequency of stochastic interconversion between the sorted MITF^high^ and MITF^low^ subpopulations during the drug treatment, we monitored the MITF activity within large numbers of single-cells, over a 5-day period of BRAFi treatment. As expected, the MITF^high^ cells always displayed higher activity (quantified by the GFP-reporter) than did the MITF^low^ cells (Fig. 3g), with no significant stochastic switching between the two trajectories observed.

To further confirm that the sorted cells reach their respective destination states after five days of drugging, we quantified the markers that are differentially expressed between the upper and lower paths at D5. Mining of the single-cell data sets revealed that several markers, including Slug, MITF, MART1 and PFK are differentially expressed between the two paths (negative-and positive-valued module 2) at D5 (Fig. 3h, Supplementary Figs. 9 and 13a). By analyzing the expression of these four genes in sorted MITF^high^ and MITF^low^ D0 cells after five days of treatment (Supplementary Fig. 13b), we found that their expression levels in sorted MITF^low^ cells were significantly lower than those in MITF^high^ cells after five days of treatment (Fig. 3i). These results experimentally support that, upon drug treatment, MITF^high^ and MITF^low^ cells take distinct trajectories toward drug tolerance along the upper and lower paths respectively (Supplementary Fig. 13a**, left panel**).

### MITF is the molecular driver for the two drug response trajectories

MITF is suggested to be an elicitor of intrinsic drug tolerance^62^. To investigate if MITF drives the bifurcation in drug response, we generated a M397 cell line with MITF stably knocked down. Before treatment, knockdown of MITF induced the cells to become slow-cycling with characteristic low Ki67 expression (Supplementary Fig. 14a, b), suggesting that downregulation of MITF will force these cells to transition along the lower path. Furthermore, upon five days of BRAFi treatment, MITF knockdown cells showed significantly lower levels of Slug, MITF, MART1 and PFK relative to control (Supplementary Fig. 14 c), suggesting that MITF-silenced cells did, in fact, follow a trajectory along the lower path. Thus, MITF is identified as an important molecular driver that discriminates between the two drug response trajectories we identified.

### Critical point analysis identifies central regulators along both trajectories

Surprisal analysis of the single-cell data sets indicates that both the upper and lower paths are characterized by a cell state transition in the D1-D3 time window (Fig 3a, **left panel**). A critical point analysis of the single-cell data in different regions of the FLOW-MAP can provide validation of this picture, and can also help identify the tipping points at which those cell state changes take place.^51,63,64^ Furthermore, network analysis of those tipping points can be used to identify key regulators that drive the transition from drug-naïve to drug-tolerant^12,51,58,63,65,66^.

We first clustered the single-cell data from all time points into 14 different sub-clusters on the FLOW-MAP. Clusters 1, 6, 7, 8, 10, 11 and 12 align with the upper path, while clusters 2, 3, 9, 13 and 14 fall along the lower path (Fig. 4a). Two previously reported critical state transition indices, the signaling network activity index (SNAI)^12^ and the critical transition index (Ic)^64^, were utilized to evaluate the tipping points associated with the lower and upper paths. We found cluster 7 in the upper path and cluster 9 in the lower path showed the highest values of these indices within their respective path (Fig.4 b, c **and Supplementary Figs. 15, 16, and 17)**, suggesting that clusters 7 and 9 are closest to the tipping points.

**Figure 4.**
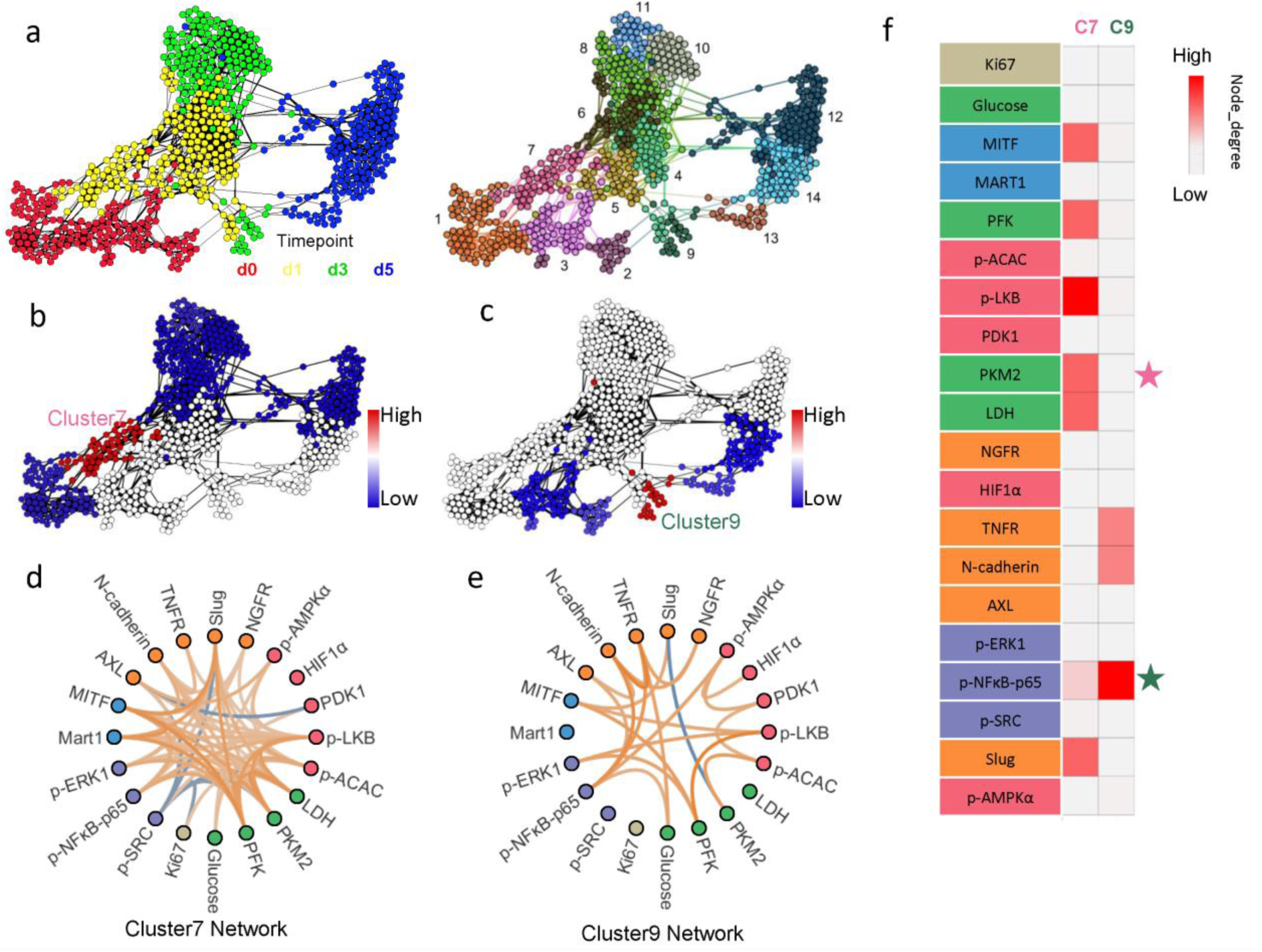
Critical point analysis (SNAI) and network analysis of two trajectories. **a.** Clustering of all cells into 4 time point-defined subpopulations. Left panel is FLOW-MAP with cells color coded by drug exposure time. Right panel is FLOW-MAP with cell color-coded as one of the 14 subpopulations defined from clustering analysis. **b.** Critical point transition analysis for upper path. Critical point index SNAI is calculated within each subpopulation associated with the upper path and color-coded onto the FLOW-MAP. Red indicates higher SNAI value, while blue represents lower SNAI value. Cluster7, shown where labeled, shows the highest SNAI value in the upper path. **c.** Critical point transition analysis for lower path. Critical point index SNAI is calculated within each subpopulation associated with the lower path and color-coded onto the FLOW-MAP. Red indicates higher SNAI value, while blue represents lower SNAI value. Cluster9, shown where labeled, shows the highest SNAI value in the lower path. **d.** Marker-marker correlation networks, extracted from SCBC data within cluster7 cells. The correlation strengths are reflected in the color of each edge (orange indicates positive correlation and blue indicates negative correlation). **e.** Marker-marker correlation networks, extracted from SCBC data within cluster9 cells. The correlation strengths are reflected in the color of each edge (orange indicates positive correlation and blue indicates negative correlation). **f.** Importance score of each node within each network, as defined by node-degree. Colors indicate the node-degree value of each node within cluster7 or cluster9 networks. Nodes labeled with stars were further-tested with drug perturbation.

We next investigated the correlation networks^12,20,44^ for clusters 7 and 9. These two networks are characterized by different structures (Fig. 4d, e), implying these transitions are regulated in different ways. We quantified the participation of each analyte (node) in the correlation networks by calculating the node degree and hub score for each node (**See Methods**). For cluster 7 (upper path), we found that several transcription factors and enzymes, including MITF, PFK, p-LKB, PKM2, LDH2 and Slug, showed high levels of network participation by both scoring metrics (Fig. 4f **and** Supplementary Fig. 18). For cluster 9 (lower path), TNFR, N-cadherin and p-NFκB-p65 appeared dominant. An interesting observation was that the markers that exhibited a high score in cluster 7 often displayed a low score in cluster 9, and vice versa, indicating that the two paths are dissimilarly regulated.

To examine if the transitions along the two paths are driven by distinct hub regulators, we perturbed the respective hub nodes identified within clusters 7 and 9, and probed for differential influence on the two trajectories. We hypothesized that inhibition of the glycolysis enzyme PKM2 and the signaling phosphoprotein p-NFκB-p65 would differentially influence the transitions along upper and lower paths respectively (Fig. 4f **and** Supplementary Fig. 18). Accordingly, we used a PKM2 inhibitor (PKM2i) or an NFκB inhibitor (NFκBi) in combination with the BRAFi to treat sorted MITF^high^ and MITF^low^ cell subpopulations. Consistent with our hypothesis, the MITF^low^ subpopulation was more sensitive to the BRAFi + NFκBi combination (Fig. 5a), while the MITF^high^ subpopulation was more sensitive to the BRAFi + PKM2i combination (Fig. 5b). This hypothesis was further validated by testing the same drug combinations on the MITF-knockdown cell line relative to unmodified M397 cells (Fig. 5c, d). Thus, cells passing along the different trajectories displayed differential sensitivities to PKM2 and NFκB inhibition.

**Figure 5.**
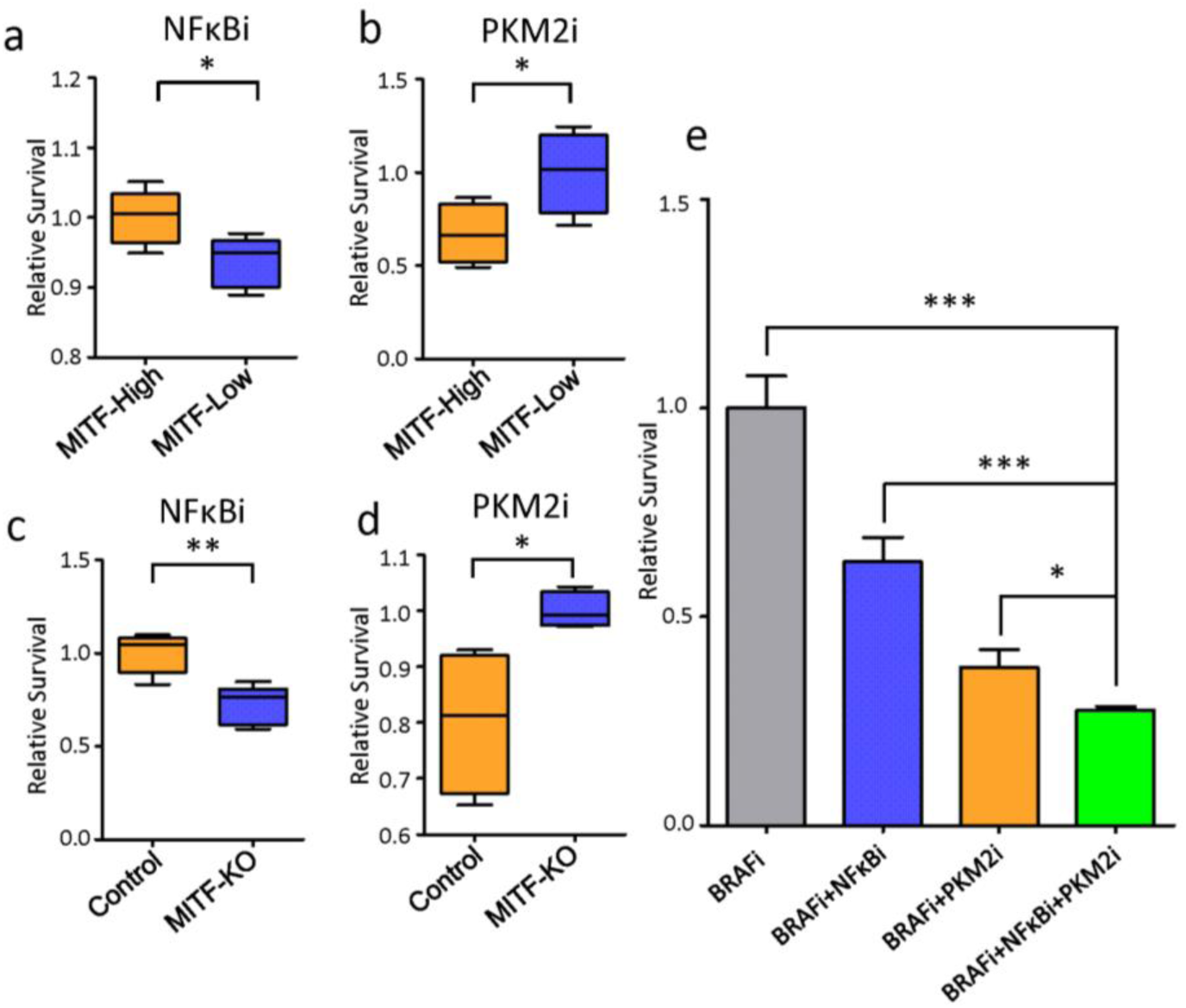
Differential drug sensitivity of cells associated with two trajectories. **a.** MITF-GFP reporter cell line were sorted for MITF-high and MITF-low subpopulation before drugging. The sorted cells were then treated with BRAFi+NFΚBi combination for 5days and then harvest for cell number counting. Relative cell survival of sorted MITF-high and MITF-low cells after undergoing BRAFi+NFΚBi combination therapy for 5 days were plotted. Survival data were normalized to MITF-high sample. **b.** MITF-GFP reporter cell line were sorted for MITF-high and MITF-low subpopulation before drugging. The sorted cells were then treated with BRAFi+PKM2i combination for 5days and then harvest for cell number counting. Relative cell survival of sorted MITF-high and MITF-low cells after undergoing BRAFi+PKM2i combination therapy for 5 days were plotted. Survival data were normalized to MITF-low sample. **c.** MITF knockdown cells and control cells were treated with BRAFi+NFΚBi combination for 5days and then harvest for cell number counting. Relative cell survival of sorted control and MITF-sh cells after undergoing BRAFi+NFΚBi combination therapy for 5 days were plotted. Survival data were normalized to control sample. **d.** MITF knockdown cells and control cells were treated with BRAFi+PKM2i combination for 5days and then harvest for cell number counting. Relative cell survival of sorted control and MITF-sh cells after undergoing BRAFi+PKM2i combination therapy for 5 days were plotted. Survival data were normalized to MITF-KO sample. **e.** M397 cell treated with BRAFi, BRAFi+NFΚBi, BRAFi+PKM2i and BRAFi+NFΚBi+PKM2i for 5 days were harvest for cell number counting. Relative cell survival of cells after undergoing BRAFi, BRAFi+NFΚBi, BRAFi+PKM2i, or BRAFi+PKM2i+NFΚBi therapy for 5 days were plotted. Survival data were normalized to cells undergoing BRAFi monotherapy treatment.

Considering the differential regulator dependence of the two trajectories, we further hypothesized that co-blocking both trajectories by simultaneously inhibiting PKM2 and NFκB signaling might show additive effects in preventing the transitions towards BRAFi tolerance. To test this hypothesis, we used the triple drug combination (BRAFi + PKM2i + NFκBi) to treat the M397 cells *in vitro* for five days and compared the resulting cell number against monotherapies (BRAFi only) and double-drug combinations (BRAFi + PKM2i and BRAFi + NFκBi) for five days. Consistent with our prediction, the triple-drug combination significantly outperformed the double-drug combinations which in turn were superior to the monotherapy (Fig. 5e). Further, PKM2i or NFκBi monotherapy showed minimal growth inhibition on the M397 cells (Supplementary Fig.19), implying that these drugs likely function by selectively blocking the BRAFi-induced cell state transitions to the drug-tolerant state. These results demonstrate that the upper and lower paths are independent, have different regulators, and are independently druggable.

## Discussion

We explored here whether cell trajectories connecting between the initial and final states of a cell-state transition could be determined from a kinetic series of static snapshots of the traversed cell-state space landscape. As a model system, we utilized a highly plastic, patient-derived M397 *BRAF^V600E^* mutant melanoma cell line, which has been shown to reversibly transition between drug-naïve and drug-resistant states upon treatment with a BRAF inhibitor. While single-cell omics tools have proven immensely valuable for resolving the cellular heterogeneity of tissues at a single given time point, here we sought to quantitatively connect that cellular heterogeneity to dynamic heterogeneity of cell state changes.

We utilized microfluidic-based SCBC technology to characterize the cellular heterogeneity during the first five days of drug-response. Because both metabolic activity and signaling pathways display functional changes during the early drug-response, SCBC is uniquely suited here since it is capable of simultaneously capturing both metabolites and cytoplasmic proteins (and phosphoproteins) from single cells. However, unlike single-cell RNA-seq, single cell proteomics is typically limited to assaying only tens of functional proteins and metabolites. In order to accurately capture the cell state space accessed by M397 cells under BRAFi treatment, we first utilized transcriptomic analysis and literature guidance to define a panel of 20 analytes that included phenotypic markers, and markers of metabolic activity, oncogenic signaling, and cell proliferation, all of which are altered during the initial drug-response. Single cell analysis using this carefully selected panel readily resolved the complex cell-state space traversed by the cells during the first few days of BRAFi treatment. Of course, moving towards larger numbers of analytes would certainly provide for a deeper characterization.^67–69^

We utilized computational and theoretical methods^32,33,36,37,70–73^, integrated with additional cell biology experiments, to translate the SCBC kinetic series of snapshots in to classes of single cell trajectories. Dimensional reduction of the dataset using the FLOW-MAP algorithm revealed suggested that the cells might take one of two paths (labeled “upper” and “lower”) through cell-state space that connected the drug-naïve cells to the drug-resistant cells. Surprisal analysis of the same data resolved both a time-dependent module and a path-dependent module. The path-dependent module suggested that cells traveling along one path are separated from the other path by a biophysical barrier, which appeared to be associated with the transcription factor MITF and its downstream melanocytic marker MART1. These analyses further predicted that the trajectory a specific cell takes is determined by its MITF level prior to drug treatment. These predictions were verified experimentally, which supported the integration of computational visualization methods with theoretical biophysical approaches to gain insight into a complex biological system. Such an approach should be broadly applicable to other dynamic, complex biological systems, including studies of cellular differentiation, tumorigenesis, and more.

Proliferative and invasive phenotypes are well-known in melanoma^61,74^. MITF, MART1 and Ki67 have been reported as robust markers for distinguishing these two phenotypes^61,74^. We have found that these two distinct phenotypes can co-exist even in the untreated, isogenic M397 cell line used in our study. The MITF^high^ and MITF^low^ subpopulations not only displayed different doubling time without BRAFi treatment but also followed distinct drug-response trajectories upon treatment. This finding is consistent with the observations of melanoma phenotype switching from a melanocytic and highly proliferative state to a non-melanocytic and more invasive state^61^. In that study, proliferative or invasive cell lines displayed fixed gene expression profiles in culture, but when transplanted *in vivo*, each class generated heterogeneous tumors containing cells with both kinds of expression profile. Consistent with their observation of fixed gene expression profiles *in vitro*, we did not observe significant inter-conversion between cells traveling along different paths during the five-day treatment period. These findings suggested that these two phenotypes are relatively stable in short term period of BRAFi treatment *in vitro*. Of course, our *in vitro* study may not fully recapitulate *in vivo* melanoma biology in which the tumor microenvironment can wield a strong influence. Furthermore, we also found that transition towards MITF-low invasive-like phenotype can be easily induced by artificial knockdown of a single transcription factor: MITF. This indicates that the complex cell-state landscape is likely regulated by very few master-regulators. It also emphasizes the importance of MITF as a molecular driver in regulating melanoma phenotype determination^75^. These findings, which add significantly to our understanding of melanoma phenotype regulation, would not have been evident had it not been for single-cell analytics.

The coexistence of two distinct drug-response trajectories even in an isogenic cell line may explain the so-called “mixed-responses”, which is commonly observed during the therapeutic treatment of melanoma in clinical settings. Such alternative “escape paths” may also explain why melanomas are so refractory to BRAFi targeted therapy. Intriguingly, for each of the two paths, different drug-susceptibilities were identified by critical point analysis and network analysis: the upper path was found to be susceptible to inhibition of the glycolysis enzyme PKM2, while the lower path is sensitive to NFκb-p65 inhibition. These differential drug sensitivity results are consistent with previous bulk studies on invasive phenotypes of melanoma: MITF-low, invasive (or mesenchymal) melanoma cells have been reported to be more dependent on NFκB signaling^12,76^, and the single-cell resolution of our study reveals the exact molecular and cellular dynamics behind that observation. Co-inhibition of PKM2 and NFκB pathways demonstrated superior effects in inhibiting tumor growth, however, both genes are essential regulators in normal cells and their inhibition can cause toxicity to non-malignant tissue^77,78^. Nevertheless, the resolved heterogeneous drug response trajectories update the current understanding of resistance development, and can provide a powerful methodology for identifying effective therapy combinations.

## Methods

### Cell lines, reagents and cell culture

Patient-derived melanoma cell line, M397, used in this study was generated under UCLA IRB approval # 11–003254. Cells were cultured at 37 °C with 5% CO2 in RPMI 1640 with L-glutamine (Life Technologies), supplemented with 10% fetal bovine serum (Omega), and 0.2% antibiotics (MycoZapTM Plus-CL from Lonza). The cell line was periodically authenticated to its early passage using GenePrint® 10 System (Promega). Presence of mutations in the genes of interest was checked by Onco Map 3 or Iontrone, and was confirmed by PCR and Sanger sequencing as previously described^79,80^. BRAF inhibitor (vemurafenib), PKM2 inhibitor (Compound 3K) and NFκB inhibitor (JSH-23), all from Selleck Chemicals LLC, were dissolved in DMSO at designated concentrations before applying to cell culture media. M397 cells were plated in 10cm tissue culture plate at 60% confluency and treated with 3 µM BRAF inhibitor for the specified numbers of days.

### Microchip fabrication and integrated single-cell proteomic and metabolic assay

The fabrication of the SCBC devices and the protocol of the integrated single-cell proteomic assays were extensively discussed in our previous publications^44,46^. Briefly, the DNA microarrays within each microchamber were converted to antibody or Nano-probe microarrays by flowing the DNA-antibody or DNA-probe conjugate cocktail solution immediately before use. Cells treated with Gluc-Bio^46^ were randomly loaded into microchambers within the SCBC. Each microchamber has an assay component, and a separate reservoir of lysis buffer, and was photographed after cell loading. The SCBC was then cooled on ice for cell lysis. Following a 2-hour protein and metabolite capture period at room temperature, the microchambers were flushed and the captured protein or metabolite on the arrays were converted into fluorescent readout and digitized by a Genepix scanner (Molecular Devices).

### Data processing from Genepix scanner

By a custom MATLAB code, the average fluorescence signals for all bars within a given barcode were extracted and matched with the micrograph of that array to prepare a table that contains the microchamber address, the numbers of cells, and the measured fluorescence levels of each assayed protein or metabolite. The SCBC readouts from the microchambers with a single cell were collected to form an m × n matrix table where each row (m) represents a specific microchamber address and each column (n) represents the abundance of a specific analyte. This matrix table is used for further analysis.

### FLOW-MAP Visualizations

All FLOW-MAP visualizations were created with the FLOWMAPR R package available on GitHub (https://github.com/zunderlab/FLOWMAP/). Graphs were produced with seed.X = 1 and no clustering or downsampling. Final figures were produced in Gephi (https://gephi.org/) either using the “bluered” palette described in the FLOWMAPR package or using the “jet” rainbow palette. The code used to generate the exact FLOW-MAP graphs is available upon request.

### Surprisal Analysis

Surprisal analysis was applied as previously described^57^. Briefly, the measured level of analyte i at cell c, ln *X*_*i*_(*c*), is expressed as a sum of a steady state term ln 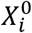(*c*), and several constraints (modules) *λ*_*j*_(*c*) × *G*_*ij*_ representing deviations from the steady state. Each deviation term is a product of a cell-dependent weight (influence score) of the constraint *λ*_*j*_(*c*), and the cell-independent contribution of the analyte to that constraint (module) *G*_*ij*_. To implement surprisal analysis, we compute the singular value decomposition (SVD) of the matrix ln *X*_*i*_(*c*). This factors this matrix in a way that determines the two sets of parameters that are needed in surprisal analysis: the Lagrange multipliers (*λ*_*j*_) for all constraints (modules) at a given time point, and for all times and the *G*_*ij*_ (time-independent) analyte patterns for all analyte i at each constraint j. In figure 3, cells with the top 10% most positive module2 score are defined as Module2-High cells (M2-High cells), and the most negative 10% ones are defined as Module2-Low cells (M2-Low cells).

### Time-lapse microscopy

Movies were acquired on an Olympus IX8 inverted fluorescence microscope with hardware autofocus (ZDC2) and an environmental chamber maintaining a 37C, 5% CO2 culture environment. Automated acquisition software (METAMORPH, Molecular Devices) was used to acquire differential interference contrast (DIC) and GFP images every 15 min from multiple stage positions.

### Image segmentation and single-cell fluorescence calculation

Custom MATLAB code (R2017a, MathWorks) was used to pre-process the DIC images of each movie. DIC images were first corrected for uneven illuminations of the field, then adjusted contrast to sharpen the cell edges. The processed DIC images were then segmented using image segmentation software ilastik^81^ (version 1.3.2) to acquire segmented cell bodies. 6 frames (out of 474 frames) were used as the training set for image segmentation of each movie. Pixel Classification feature of ilastik 1.3.2 was used to segment pixels of all 474 frames into ‘Background’, ‘Cell edge’ and ‘Cell body’ based on the labeled 6-image training set of each movie. GFP fluorescence data was extracted from cell body segments using a custom Python code. In each movie frame, each separated ‘Cell Body’ pixel block from DIC segmentation was first labeled as separated individual single cell. Then GFP fluorescence of each single cell block was calculated by integrating fluorescence from the corresponding pixels from GFP images. Background GFP fluorescence was calculated by the median GFP values of ‘Background’ pixels, and was subtracted from GFP values of ‘Cell Body’ pixels. Mean and standard error of the mean (SEM) were calculated for each time point from ensemble single-cell GFP fluorescence.

### Single-Cell Clustering

Prior to clustering, all single-cell data were separated by time point (i.e. day 0, day 1, day 3, and day 5). Rclusterpp clusters then applied which cluster the cells into 14 subpopulations. Rclusterpp clusters were produced using the Rclusterpp R package using all default settings (https://github.com/nolanlab/Rclusterpp). All clustering algorithms were performed with cells clustered on the following markers: Ki67, Mart1, HIF1a, LDH, AMPKA, p-ERK1, PFK, p-ACAC, Slug, and p-LKB. The code used for clustering is available upon request.

### Signaling Activity Indices

The signaling network activity index (SNAI) value is defined as “the reciprocal of the determinant of the protein-protein correlations” in previous publications^12^. The Ic value or critical transition index is defined as “the ratio of the average of all pairs of protein-to-protein correlation coefficients to the average of all pairs of cell-to-cell correlation coefficients” and was calculated as described previously^64^. The code used to calculate the SNAI/Ic indices for individual cell clusters is available upon request.

### Network Analysis

Pair-wise correlation matrices were calculated on within each of the 14 clusters using the Hmisc R package (https://cran.r-project.org/web/packages/Hmisc/index.html). Spearman correlations were calculated. Correlation output from the Hmisc package produces the pair-wise correlation values matrix. Bonferroni corrected p-value was used to filter the correlation network through statistical significance, and the correlation networks were drawn using a custom MATLAB code. Hub score and node degree for each marker in each correlation network were calculated using the igraph R package. Both scores were rescaled from 0 to 1 for each marker for side-by-side comparison and plotted to visualize marker-to-marker variation in hub behavior between methods of calculating correlation. The code used to perform the correlation network analysis is available upon request.

### mRNA extraction and qPCR

RNA was extracted from cells using the RNeasy Mini Kit or RNeasy plus Micro Kit (Qiagen) according to the manufacturer’s protocol. First-strand cDNA was synthesized from extracted total RNAs using the iScript cDNA Synthesis Kit (Bio-Rad). The expression of human Slug, MITF, MART1and PFK transcripts were analyzed by SYBR Green–based real-time quantitative RT-PCR (qRT-PCR) using specific primers purchased from Santa Cruz. Data were normalized to the expression of RPL19 and are expressed as fold changes.

### MITF knockdown cell line

Short hairpin RNA (shRNA) targeting the coding sequence of MITF and control shRNA were purchased from Santa Cruz. Lentiviruses encoding control shRNA and MITF shRNA were produced in HEK-293T cells by transient transfection of lentiviral based vectors and their packaging vectors psPAX2 and pMD2.G as previously described^82^. The virus was collected, filtered through a 0.45µm syringe filter after 48 hours and the M397 cells were spin-infected with viral supernatant supplemented with 10 µg/mL polybrene at 2,500 rpm and 30°C for 90 min. The transduced cells were selected using puromycin, starting at 3 days post-transduction.

### MITF Reporter Cell Line

The human *Tyrosinase Promoter* (TP) was subcloned from pLightSwitch Prom S700747 (SwitchGear Genomics, Carlsbad, CA) into the BamH1 and EcoRI sites of the lentiviral vector backbone, driving the expression of the Zsgreen gene. Lentivirus particles were generated as described above to stably transduced M397 cells. A clonal cell line was further generated via single cell sorting and expansion. Cells were then sorted as GFP^high^ and GFP^low^ population by BD FACSAria Fusion Cell Sorter for further treatment and analysis.

### Fluorescence microscopy

Images were acquired at 10X (Olympus, 10X FL PH, 0.3 NA) on an EVOS FL Auto Imaging System (Fisher Scientific) in YFP and differential interference contrast (DIC) channel. Light or laser intensity, exposure and gain were set to be the same between MITF^high^ well and MITF^low^ well.

### Clonogenic assay

M397 cells were plated onto six-well plates with fresh media at an optimal confluence. The media (with drug or DMSO) were replenished every two days. Upon the time of staining, 4% paraformaldehyde was applied onto colonies to fix the cells and 0.05% crystal violet solution was used for staining the colonies.

## Acknowledgments

We thank members of the Heath and Baltimore laboratories for helpful comments on the manuscript. We acknowledge the following agencies and foundations for support: NIH Grants U54 CA199090 (to J.R.H., D.B., W.W., and A.R.), U01 CA217655 (to J.R.H. and W.W.), P01 CA168585 (to A.R.), R35 CA197633 (to A.R.), and U54CA209971 (to S.K.P.); the Dr. Robert Vigen Memorial Fund, the Ressler Family Fund, and Ken and Donna Schultz (A.R.); the Jean Perkins Foundation (J.R.H.); ISB Innovator Award (Y.S.). L.R. was supported by the V Foundation-Gil Nickel Family Endowed Fellowship and a scholarship from SEOM. M.E.K. was supported by the National Cancer Institute of the National Institutes of Health under Award Number F99 CA212231 and Stanford University’s Diversifying Academia, Recruiting Excellence Fellowship. We acknowledge Rochelle Diamond and the Caltech Flow Cytometry Cell Sorting Facility for FACS analysis and advice.

## Author Contribution

J.R.H. and Y.S. conceived the study. Y.S. and G.D. designed the experiments. Y.S., G.D, H.C., R.Z, M.X., J.W., S.W., J.L., K.T., J.L., L.K., A.X., and L.R. performed the experiments. M.E.K., Y.S., W.W. and R.L. analyzed and interpreted the data. J.R. H., W.W., S.K.P., G.P.N, S.H., A.R., and M.E. provided conceptual advice on the data analysis and interpretation. Y.S., G.D., M.E.K. and J.R.H. wrote the manuscript. J.R.H. and D.B. supervised this study.

## Supplementary Information

### Supplementary Figures

**Supplementary Figure 1.**
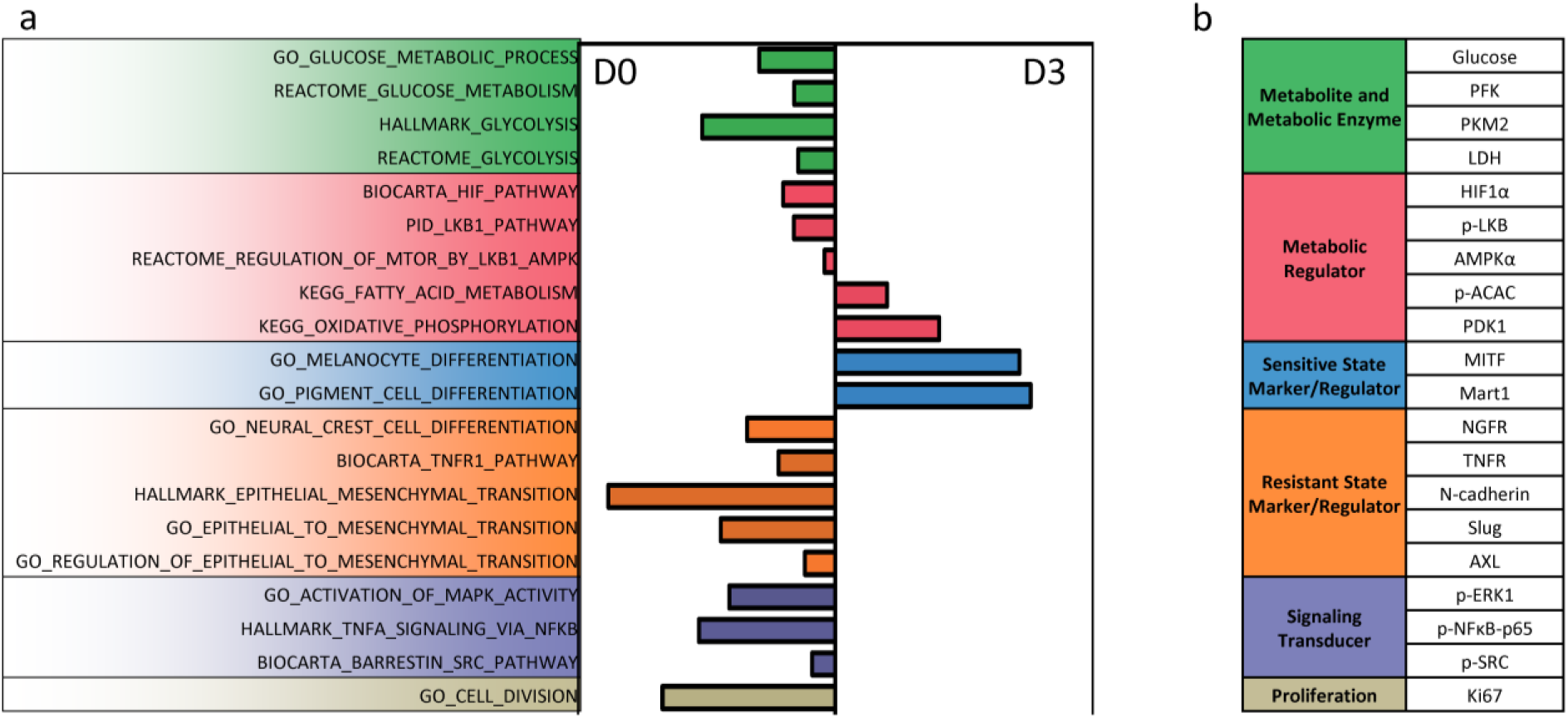
Transcriptomic analysis guided panel marker selection. **a.** Pathways that are differentially altered from day 0 to day 3 after BRAFi treatment. Each row represents a certain signaling pathway and each bar indicates normalized enrichment score (NES) calculated from geneset enrichment analysis (GSEA) of cells harvested at day 3 versus day 0. Each pathway is color-coded by its functional category as described in Fig. S1b. **b.** Panel of markers per pathway selected to quantify with single-cell barcode chip (SCBC) analysis. 20 markers were selected for SCBC analysis. Markers with similar biological function are organized together and color-coded by functional category.

**Supplementary Figure 2.**
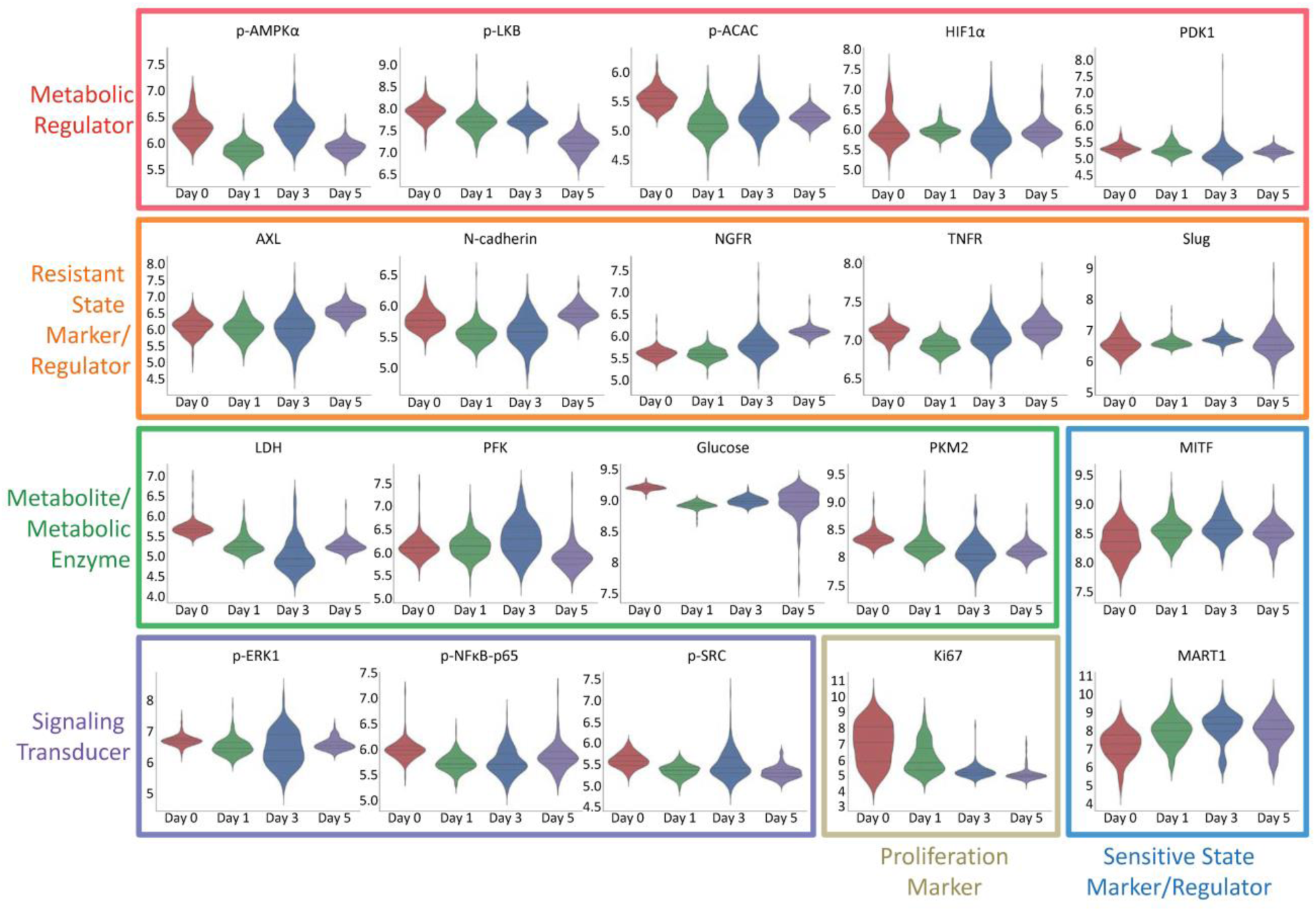
Distribution of all 20 markers across 4 time points. Each of the 20 plots represents the distributions of a certain marker level across 4 time points. Y-axis represents natural log of measured marker level. Markers within the same functional category are boxed together. Border color of each plot corresponds to the functional category of each marker, as described in Fig. 1a.

**Supplementary Figure 3.**
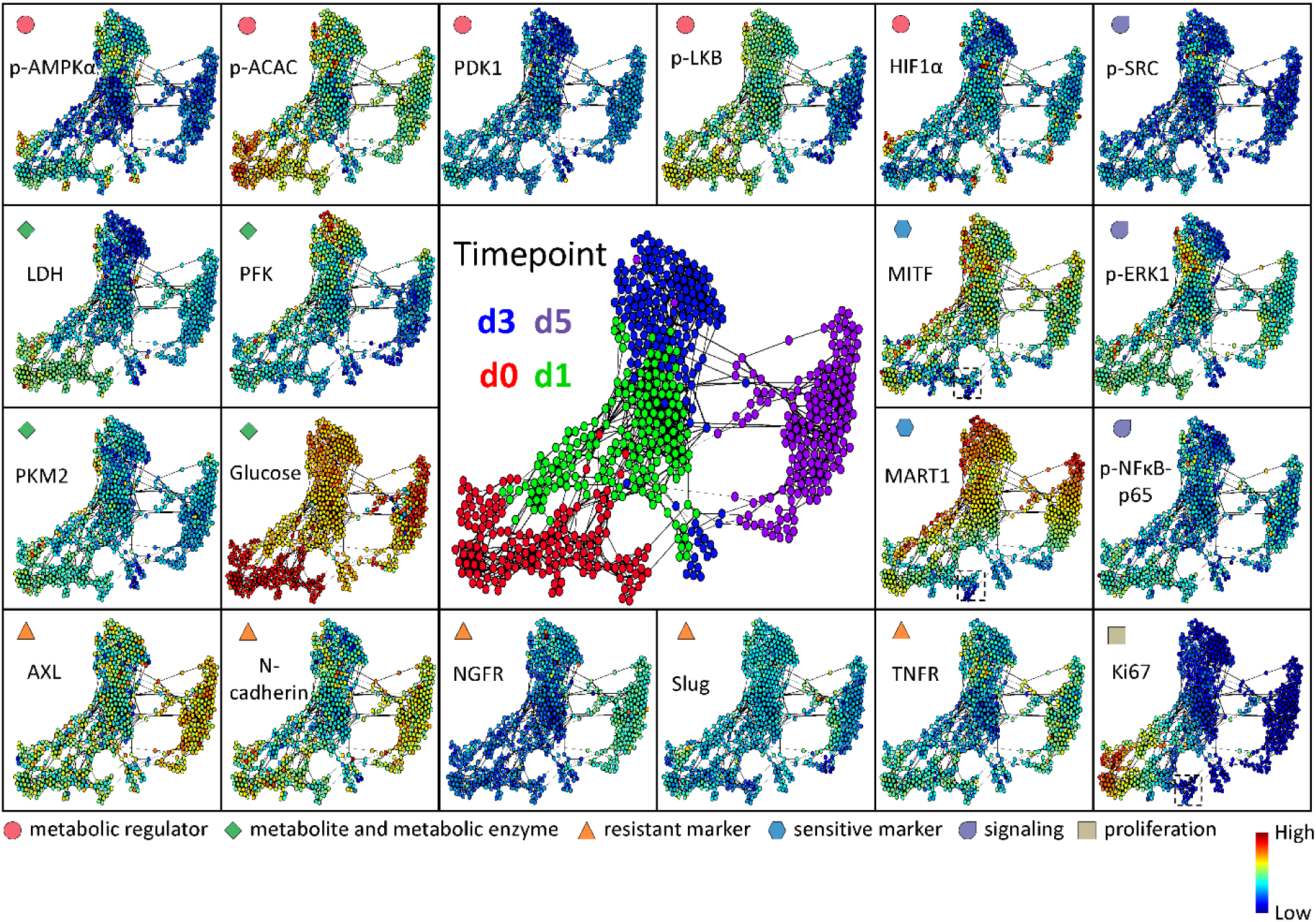
Visualization of integrated single-cell proteomic and metabolic analysis data by FLOW-MAP. Each dot represents an individual cell. The distance between each pair of cells represents the overall multi-omic dissimilarity between them. Cell pairs that are close enough are linked with an edge in between. The colors of the dots in the central panel represent BRAFi exposure time (0, 1, 3, or 5 days) of the corresponding cells. Dot colors in the other panels represent the abundance of each marker in each cell. Markers belonging to the same functional category, as described in the bottom of the figure, were assigned to a certain shape and color. The dashed-line box in the panels for MITF, MART1, and Ki67 levels shows a small subpopulation of day-0 cells that are slow cycling with less melanocytic phenotype.

**Supplementary Figure 4.**
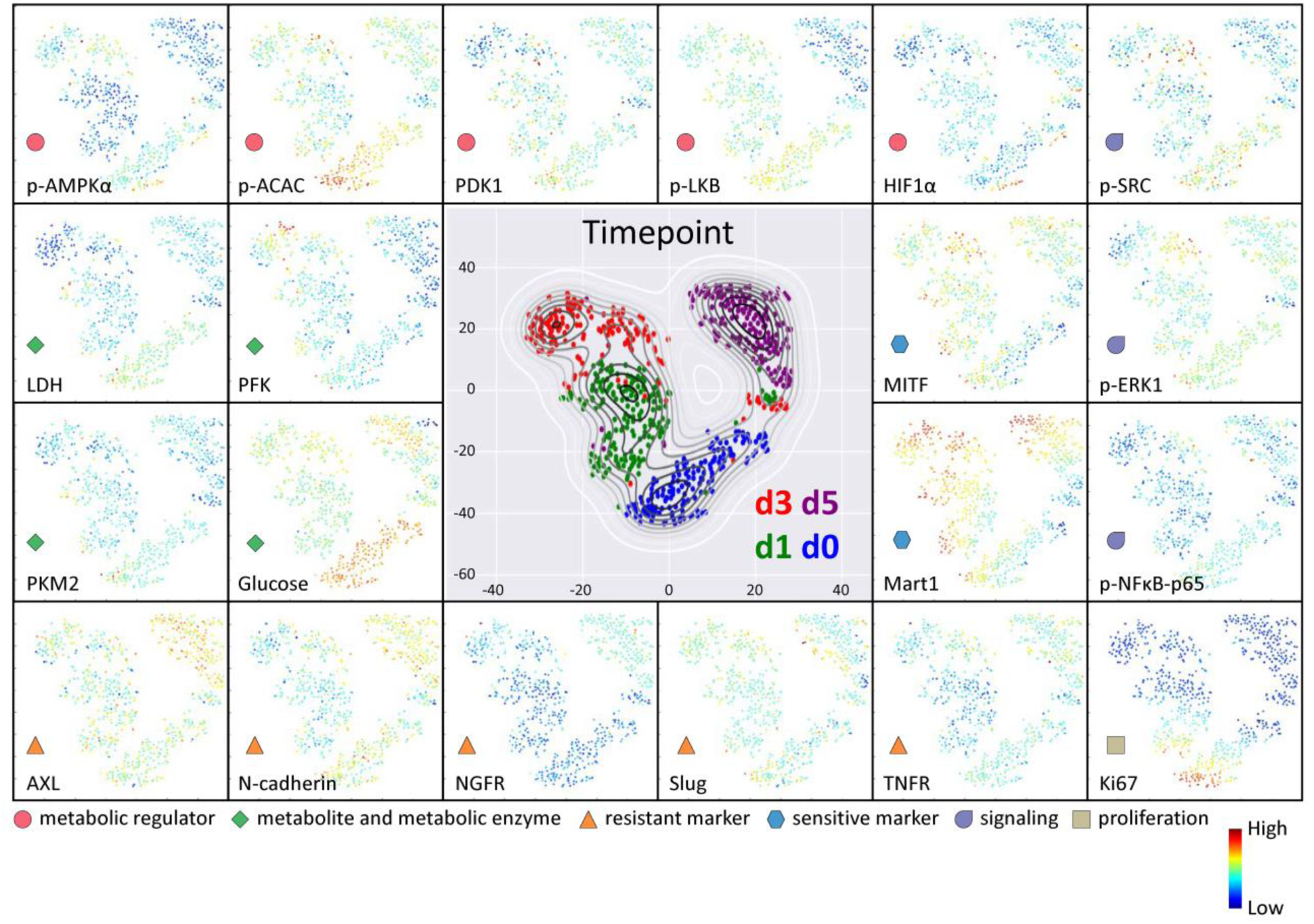
Dimension reduction with t-SNE and marker abundance visualization. Each dot per plot represents an individual cell. The distance between each pair of dots represents the overall multi-omic dissimilarity between that pair of cells. The dot colors in the central panel represent the drug exposure time of each cell. Dot colors in the other panels represent the abundance of the specified marker in each cell. Markers that belong to the same functional category were assigned to a certain shape and color, as described in the bottom of the figure. T-SNE visualizations show both the heterogeneity that exists at baseline as well as the progression across time through two separate paths.

**Supplementary Figure 5.**
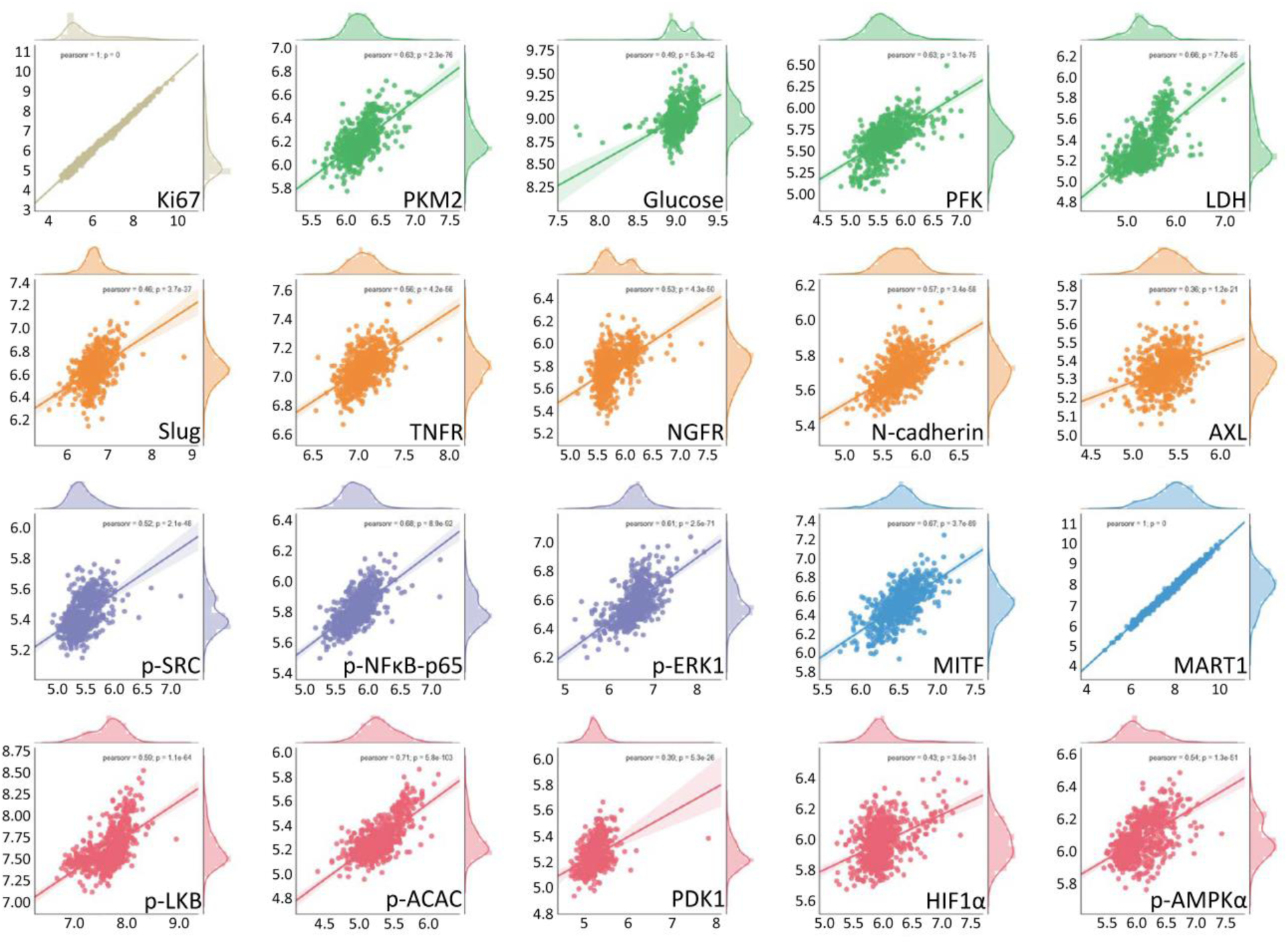
Two modules from surprisal analysis recapitulated the original experimental measured marker levels. Each plot represents an individual marker. Each dot within a single plot represents a single cell. The x-axis value of each dot represents the experimentally measured marker expression within a cell. The y-axis value of each dot represents the predicted marker level of the same cell as calculated by surprisal analysis of only module1 and module2. The strong positive correlation between the x-and y-axis values indicate that surprisal analysis of the two modules recapitulates experimentally measured marker levels per cell.

**Supplementary Figure 6.**
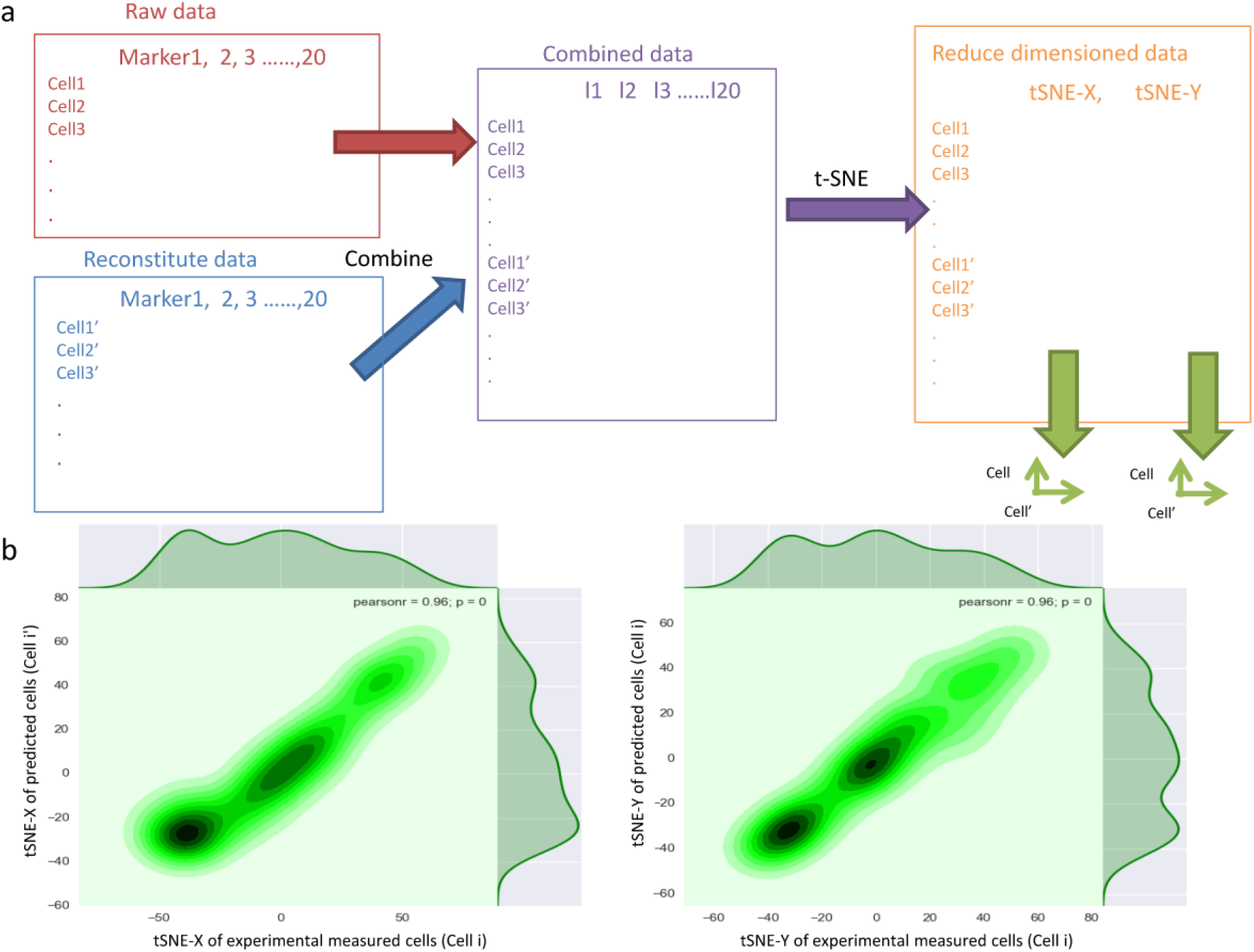
Two modules from surprisal analysis recapitulated the overall experimental measured marker levels. **a.** Schematic illustration of workflow to project raw data and surprisal analysis-predicted data onto the same 2-dimensional space. Each cell has measured levels of all 20 markers. Similarly, each cell also has predicted levels of all 20 markers as calculated from surprisal analysis. The raw and surprisal-predicted data matrices were combined to make a bigger matrix with double the original number of rows, each row representing a cell from raw data or predicted data. Each column represents a single marker, with each matrix value representing a single cell’s abundance of a marker. The combined, 20-dimensional dataset was projected onto a single t-SNE map where cells with similar levels of all 20 markers will be in nearby coordinates. **b.** Each dot represents an individual cell. In the left panel, the x-axis represents the t-SNE x-value of the cell projected from raw data, while the y-axis represents the t-SNE x-value of the cell projected from surprisal analysis-predicted data. The right panel is similar to left panel, but instead compared t-SNE y-values. The linear, x = y plots indicate that single cells, as projected from raw data and from surprisal analysis-predicted data, are in the same location in a reduced dimension; therefore, the experimentally measured and surprisal analysis-predicted expression profiles of all 20 markers are similar.

**Supplementary Figure 7.**
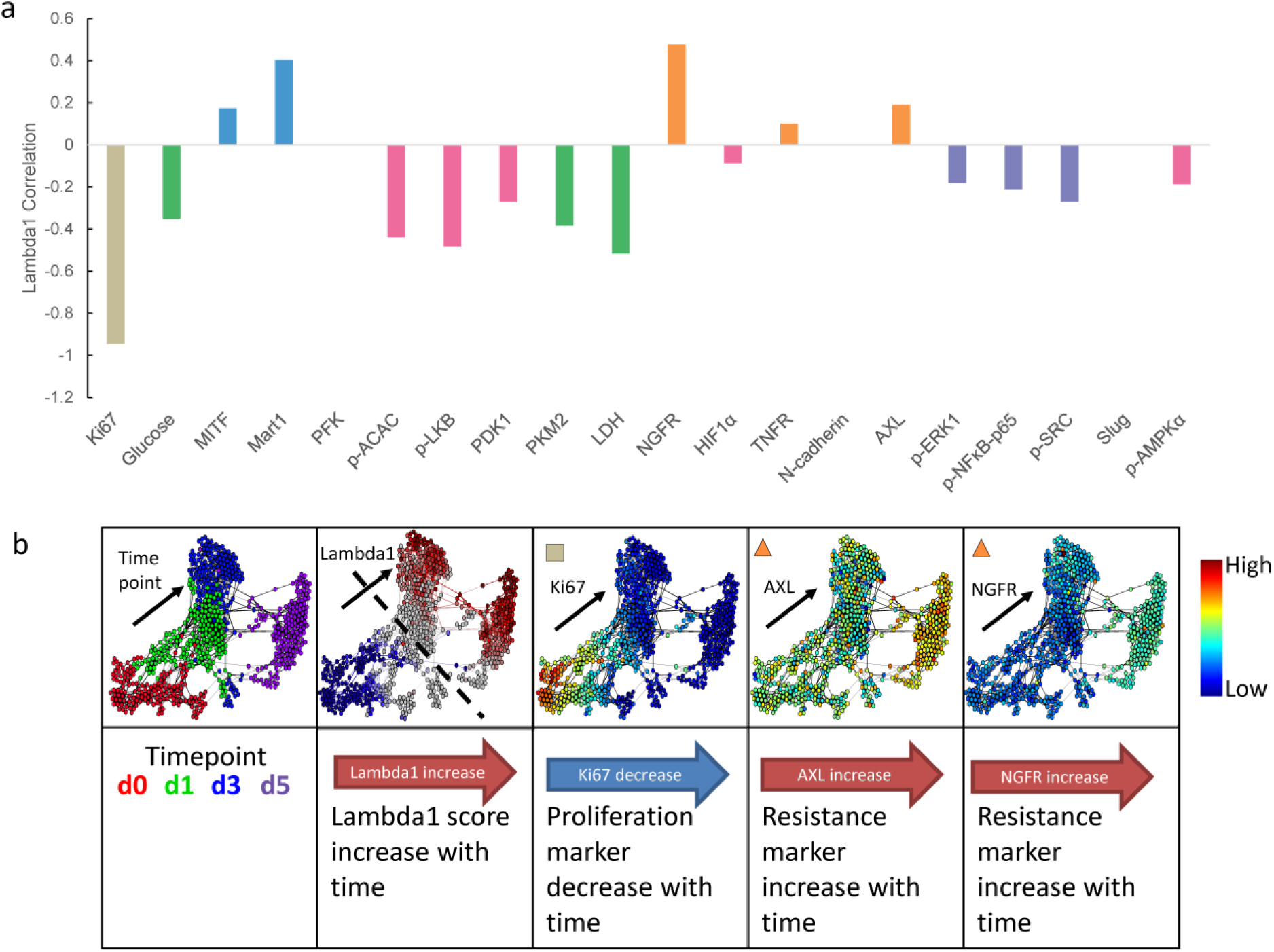
Lambda1 associated markers displayed time dependent changes. **a.** Pearson correlation of marker level vs. module1 score (lambda1) across cells from all timepoints of BRAFi exposure. Correlations that are not statistically significant (i.e. *p* > 0.05) are not shown. **b.** Representative markers that showed strongest positive (AXL, NGFR) or negative (Ki67) correlation with module1 score are shown in individual cells on FLOW-MAP.

**Supplementary Figure 8.**
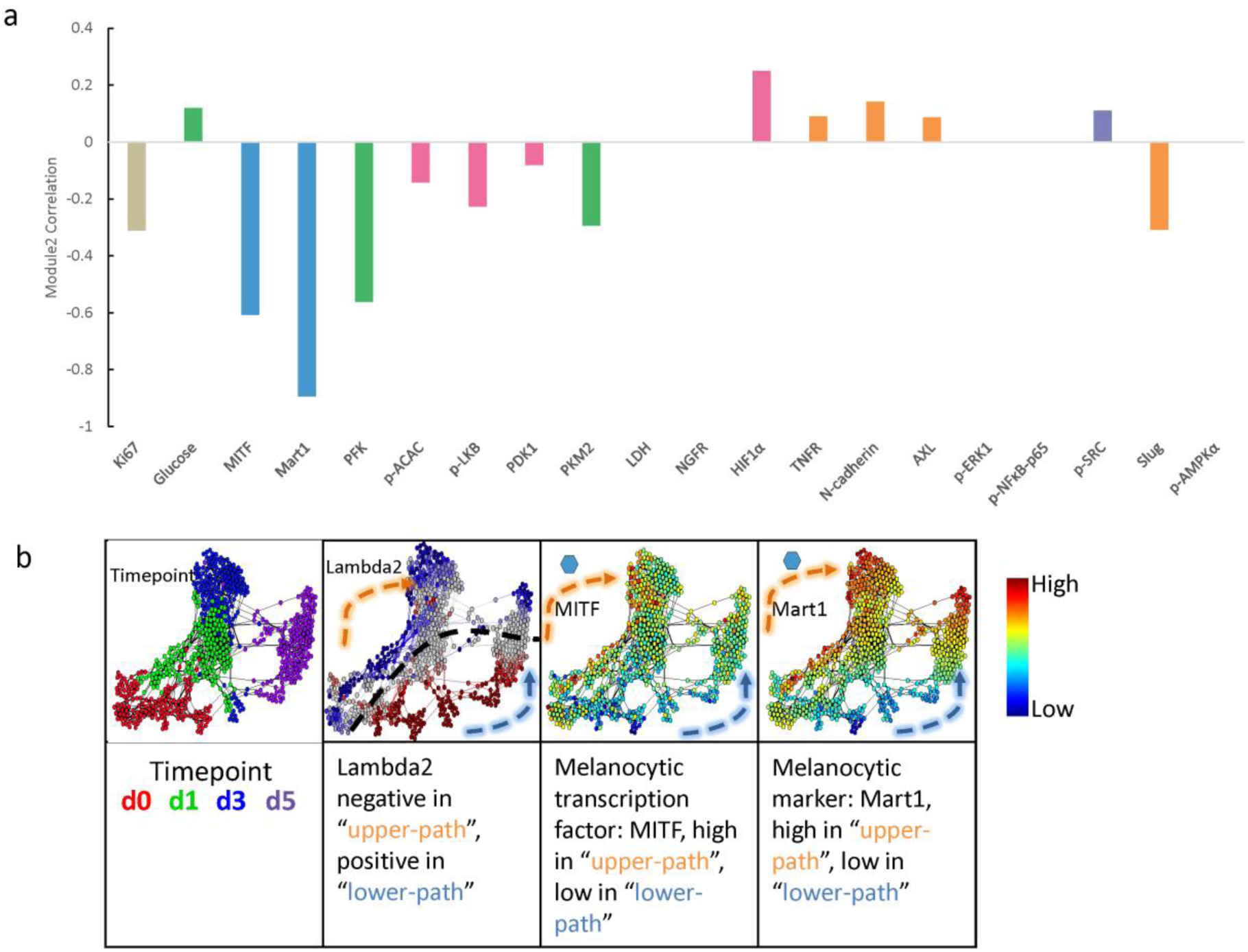
Lambda2 associated markers displayed path-specific expression patterns. **a.** Pearson correlation of marker level with module2 score (lambda2) across cells from all time points after BRAFi exposure. Correlations that are not statistically significant (i.e. *p* > 0.05) were not shown. **b.** Representative markers that showed strongest negative correlation with module1 score are shown in individual cells on FLOW-MAP.

**Supplementary Figure 9.**
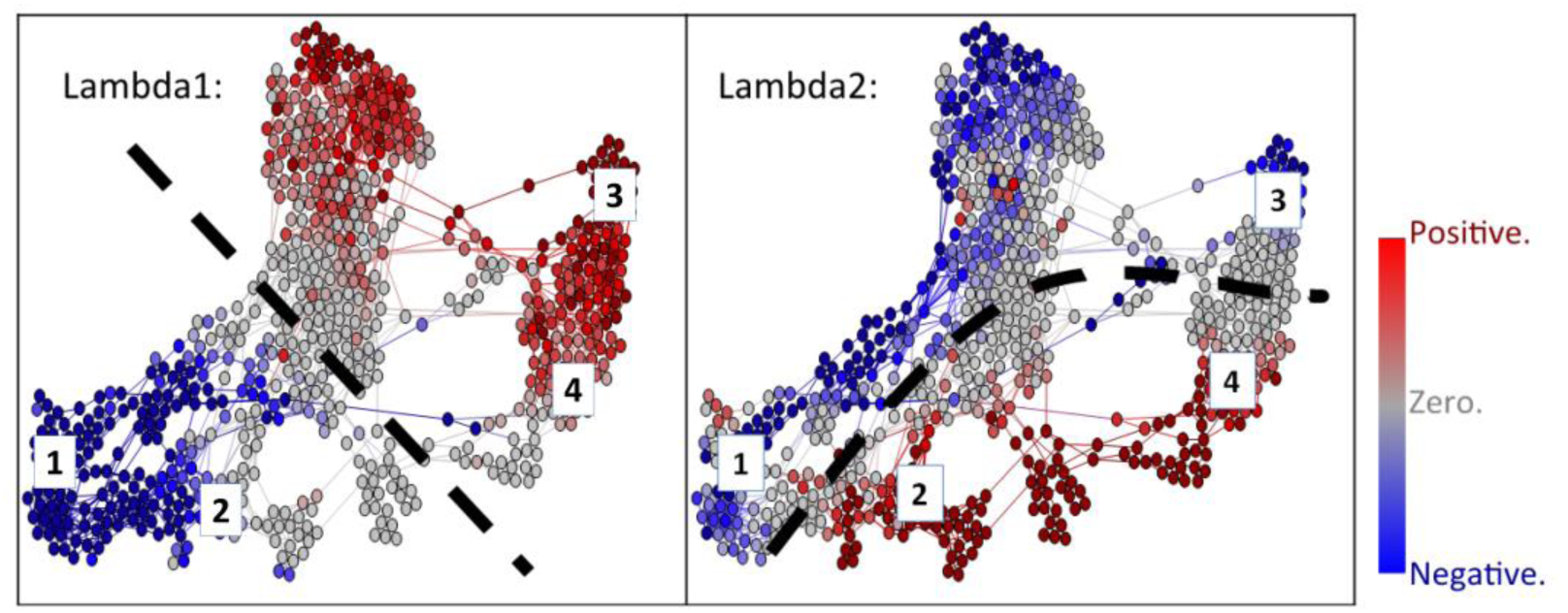
Four different cell states inferred from Module1 and Module2 associated biophysical barriers. Module1 and module2-associated barriers, as defined by the points at which a module score changes sign, separate the cells into roughly 4 different states, labeled from 1 to 4. States 1 and 2 are separated from states 3 and 4 by the module1-associated barrier. States 1 and 3 are separated from states 2 and 4 by the module2-associated barrier.

**Supplementary Figure 10.**
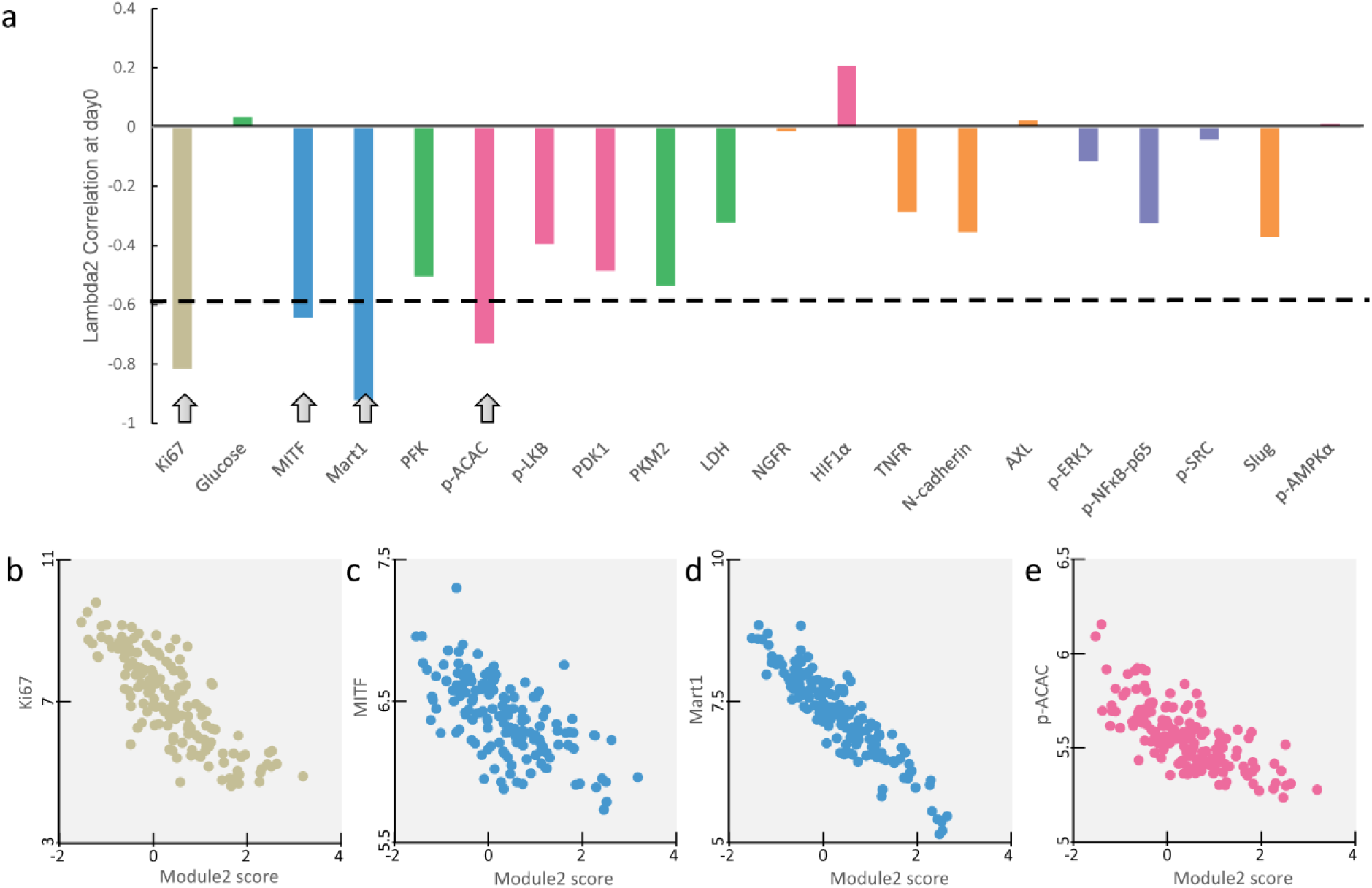
Day-0 cell analysis of marker correlation with module2, suggesting MITF as the driver for bifurcation in cell state transition trajectories. **a.** Pearson correlation of marker level and module2 score in day 0 cells from single-cell dataset. The four most highly-correlated markers are labeled with gray arrows. **b.** Scatter plots showing expression levels of the four most highly-correlated markers versus module2 score in day-0 cells from single-cell dataset.

**Supplementary Figure 11.**
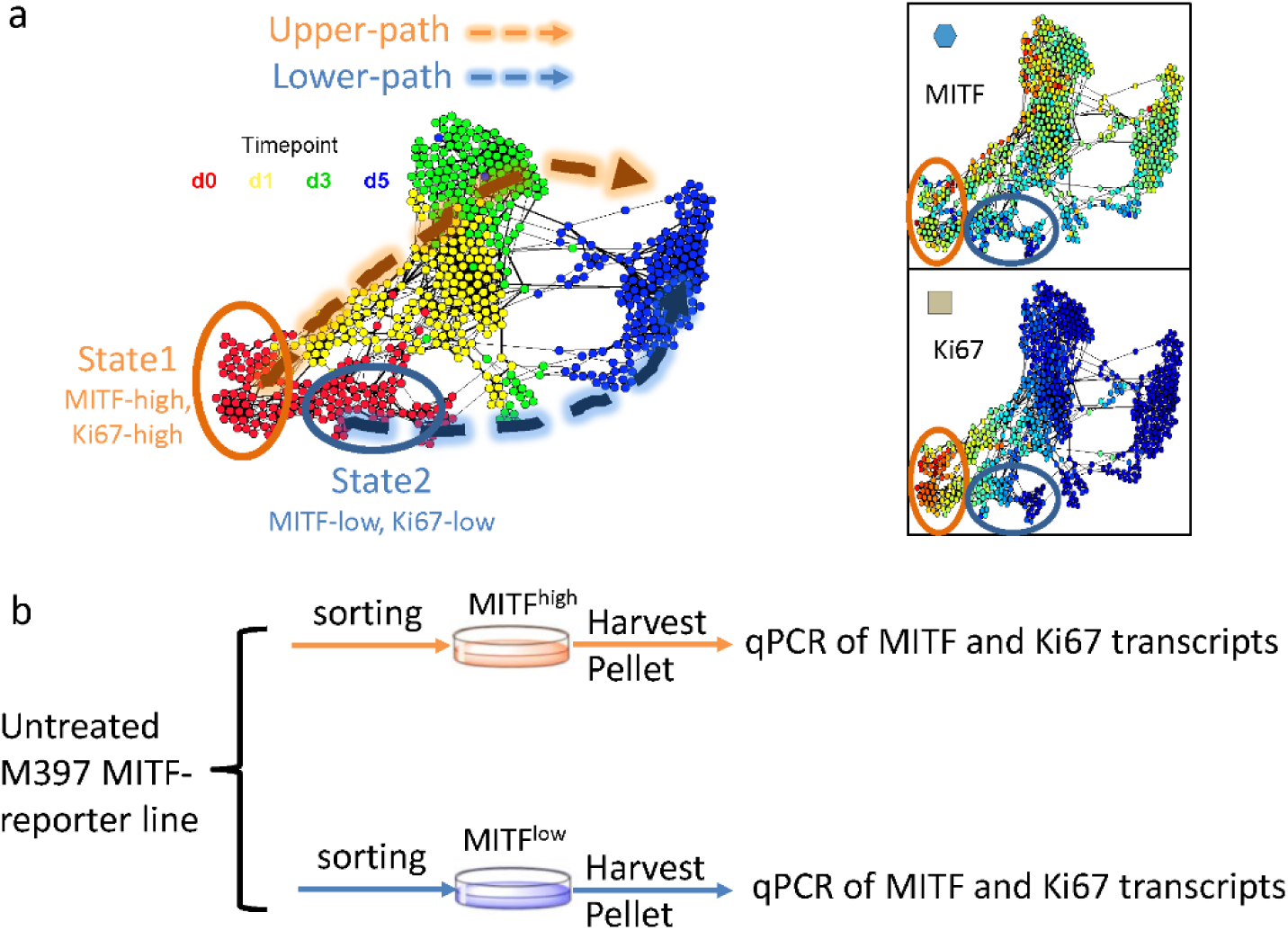
Illustration of MITF-reporter line sorting experiment on untreated cells. **a.** Untreated cells in state 1 and state 2 showed significantly different levels of MITF and Ki67. **b.** For MITF-GFP reporter line, cells with higher GFP level and lower GFP level were sorted out using FACS. The sorted cells were then harvested for qPCR quantitation of MITF and Ki67 expression.

**Supplementary Figure 12.**
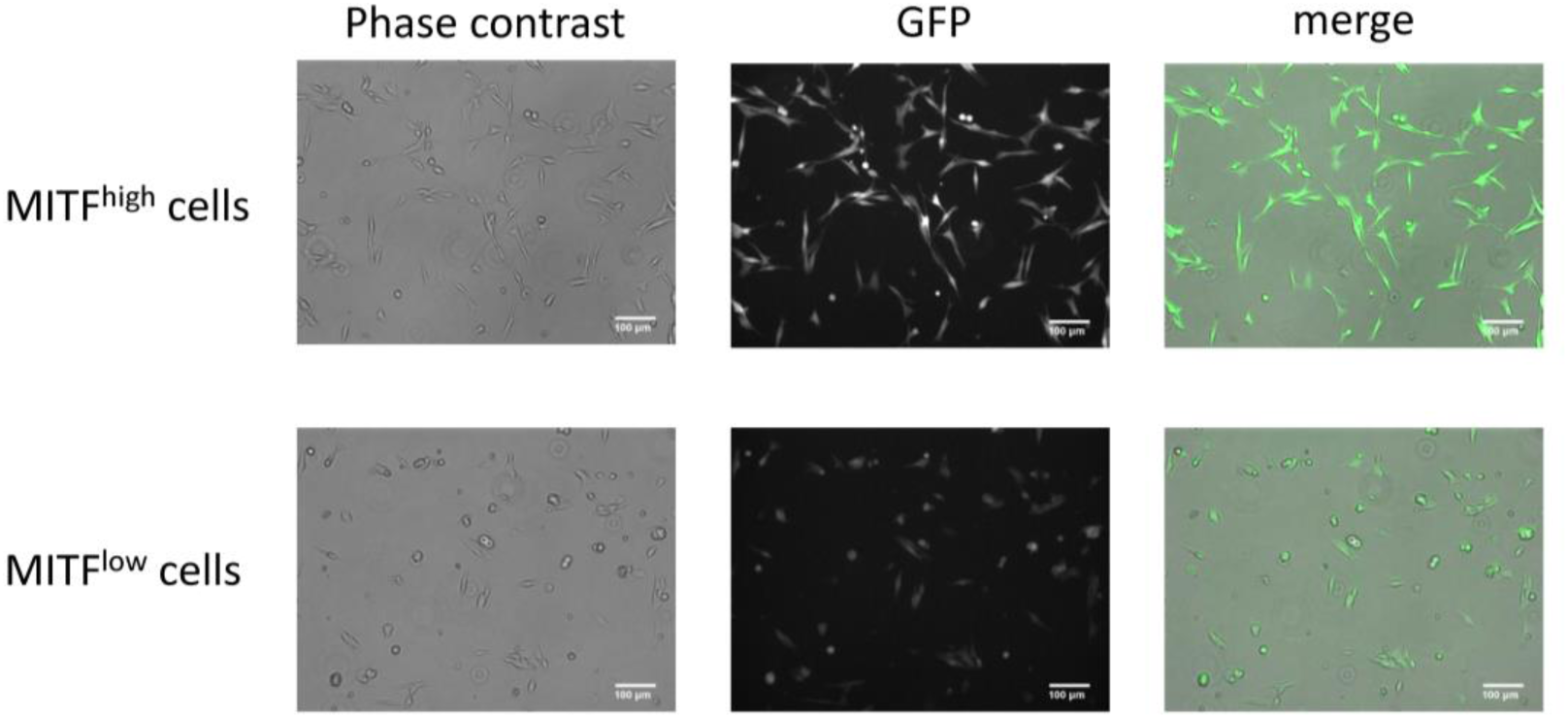
Sorted state 1 and state 2 cells shows different MITF-GFP level and morphology.

**Supplementary Figure 13.**
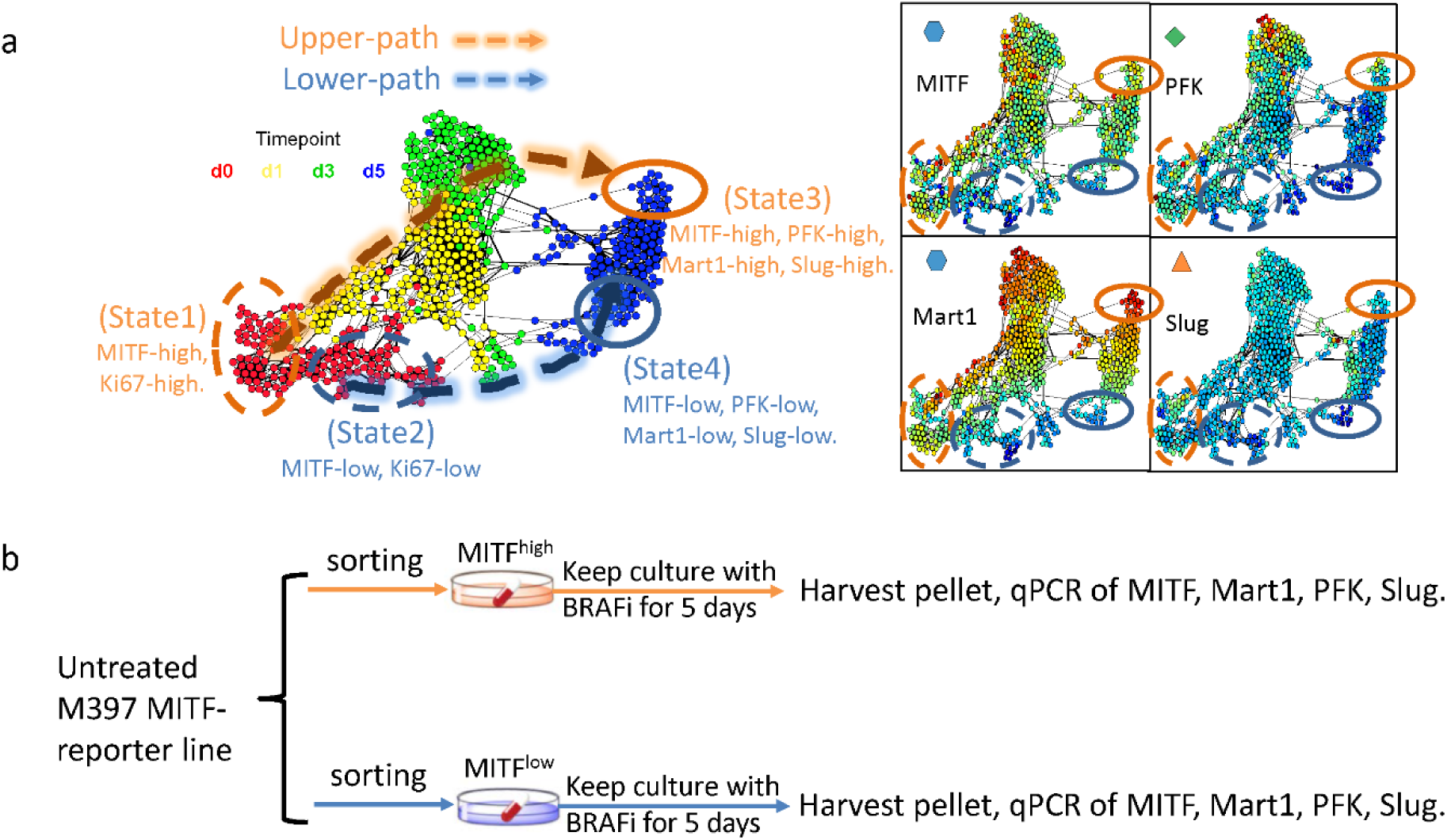
Illustration of MITF-reporter line sorting and drug treatment experiments. **a.** Day-5 cells in state 3 and state 4 showed different levels of MITF, MART1, PFK and Slug. **b.** For MITF-GFP reporter line, cells with higher GFP level and lower GFP level were sorted out using FACS. The sorted cells were then treated with BRAFi for another five days, then harvested for qPCR quantitation of MITF, MART1, PFK and Slug expression.

**Supplementary Figure 14.**
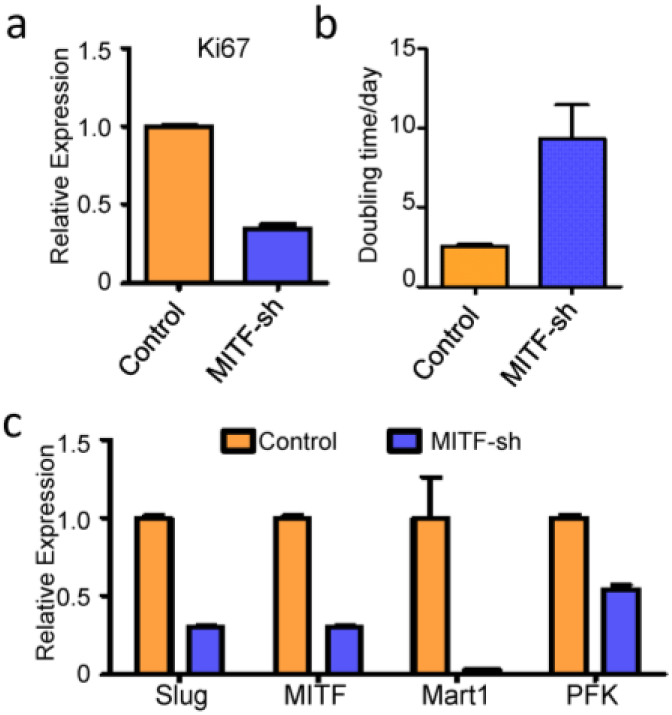
MITF knock-down cells showed similar phenotype as sorted state 2 day-0 cells which will follow the bottom trajectory to become state 4-like cells upon 5days of BRAFi. **a.** Expression level of Ki67 from qPCR of MITF knockdown cells versus control cells. **b.** Measured doubling time of MITF-knockdown cells versus control cells. **c.** Expression level of MITF, MART1, PFK and Slug after 5 days of BRAFi treatment in control cells and MITF-knockdown cells.

**Supplementary Figure 15.**
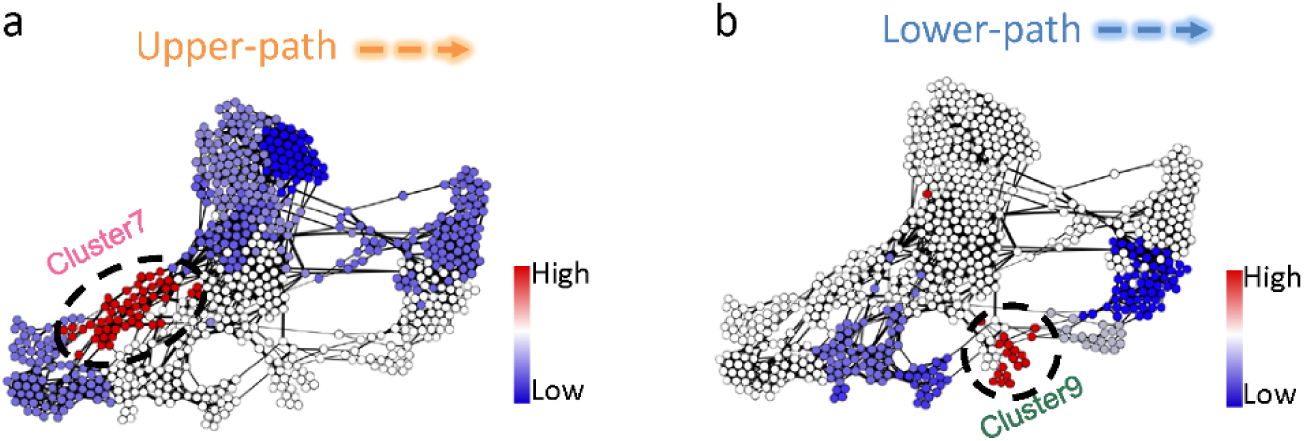
Ic value of single cells for critical point transition analysis of each trajectory. **a.** Critical point transition analysis for upper path. Critical point index Ic is calculated within each subpopulation associated with the upper path and color-coded onto the FLOW-MAP. Red indicates higher Ic value. Blue represents lower Ic value. Cluster 7, circled and labeled, shows the highest Ic value in the upper path. **b.** Critical point transition analysis for lower path. Critical point index Ic is calculated within each subpopulation associated with the lower path and color-coded onto the FLOW-MAP. Red indicates higher Ic value. Blue represents lower Ic value. Cluster 9, circled and labeled, shows the highest Ic value in the lower path.

**Supplementary Figure 16.**
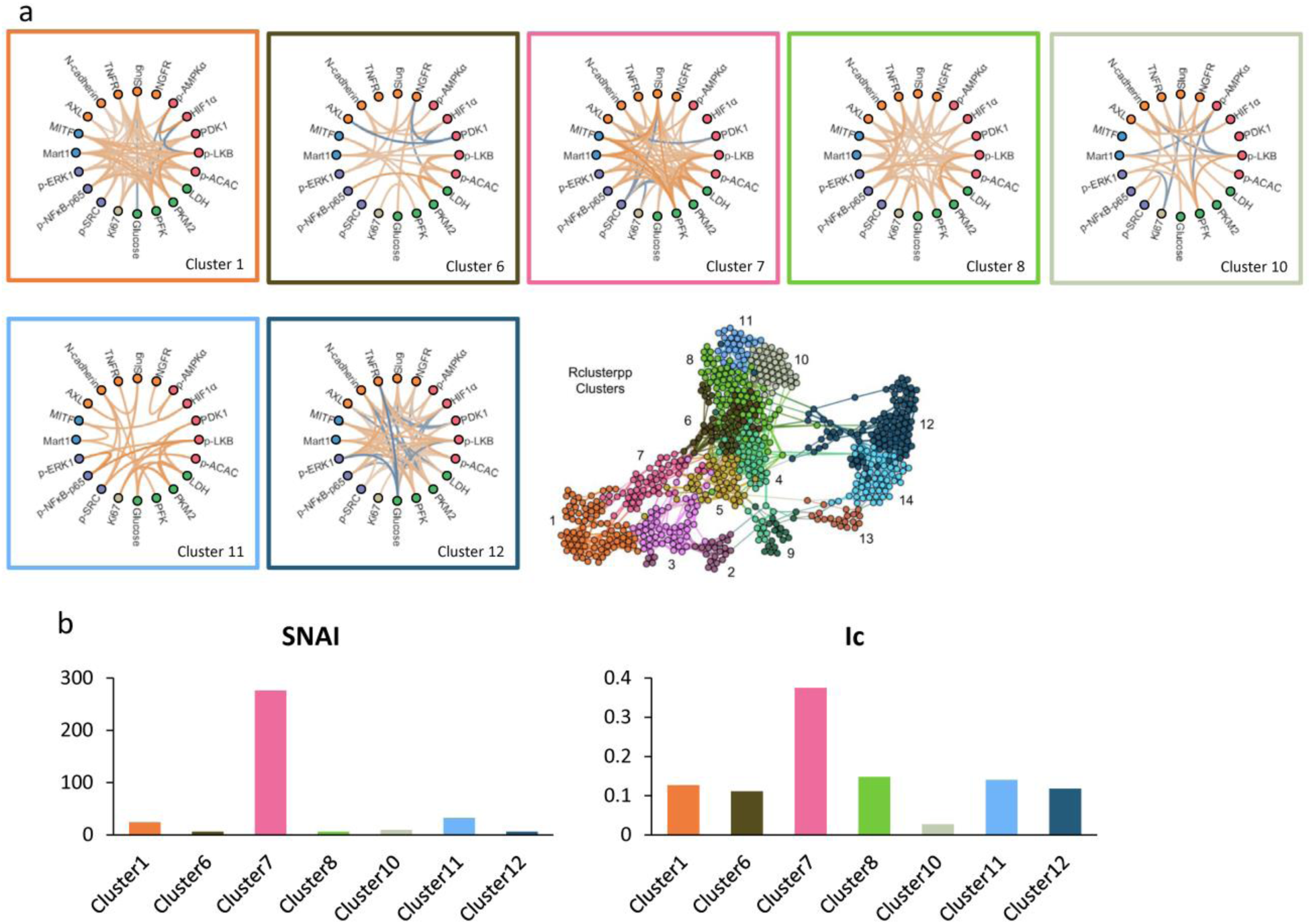
Network structure and respective SNAI and Ic values for subpopulations associated with the upper path. **a.** Network of subpopulations associated with the upper path. Each network structure plot is bordered by the color label of the corresponding cluster. **b.** SNAI and Ic values of networks associated with subpopulations in the upper path.

**Supplementary Figure 17.**
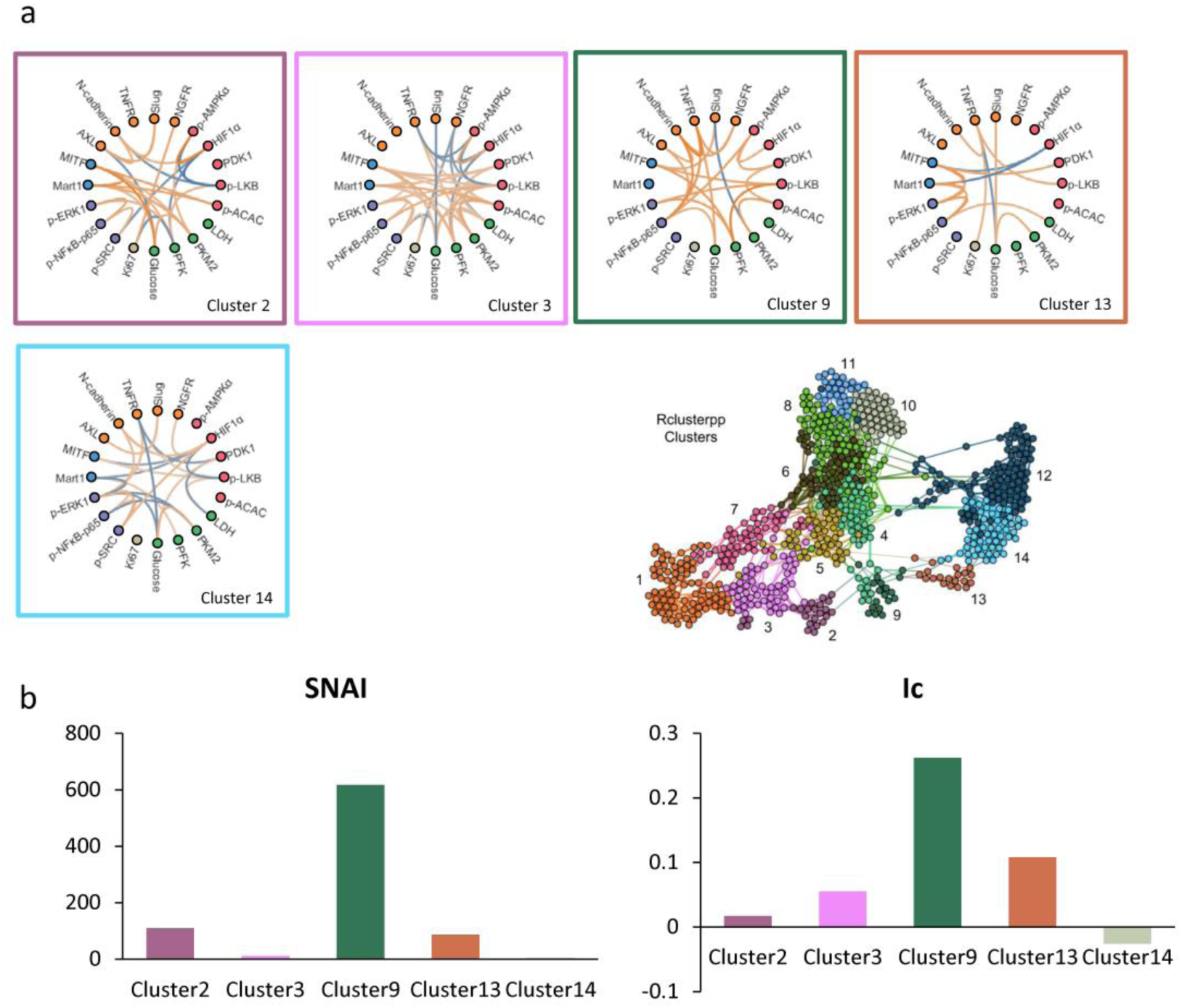
Network structure and respective SNAI and Ic values for subpopulations associated with the lower path. **a.** Network of subpopulations associated with the lower path. Each network structure plot is bordered by the color label of the corresponding cluster. **b.** SNAI and Ic values of networks associated with subpopulations in the lower path.

**Supplementary Figure 18.**
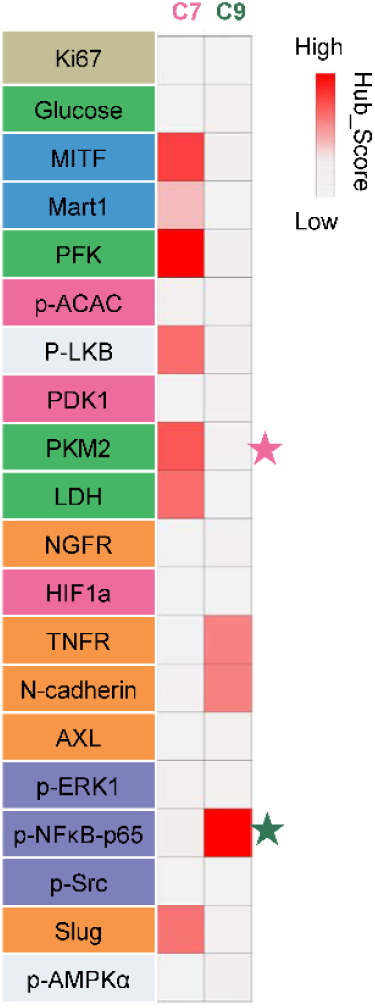
Hub-score of each node at networks for cluster7 (C7) and cluster9 (C9). Colors in C7 and C9 columns indicate the hub-score value of each node found within the cluster 7 or cluster 9 networks, respectively. Nodes labeled with stars were further tested using drug perturbation.

**Supplementary Figure 19.**
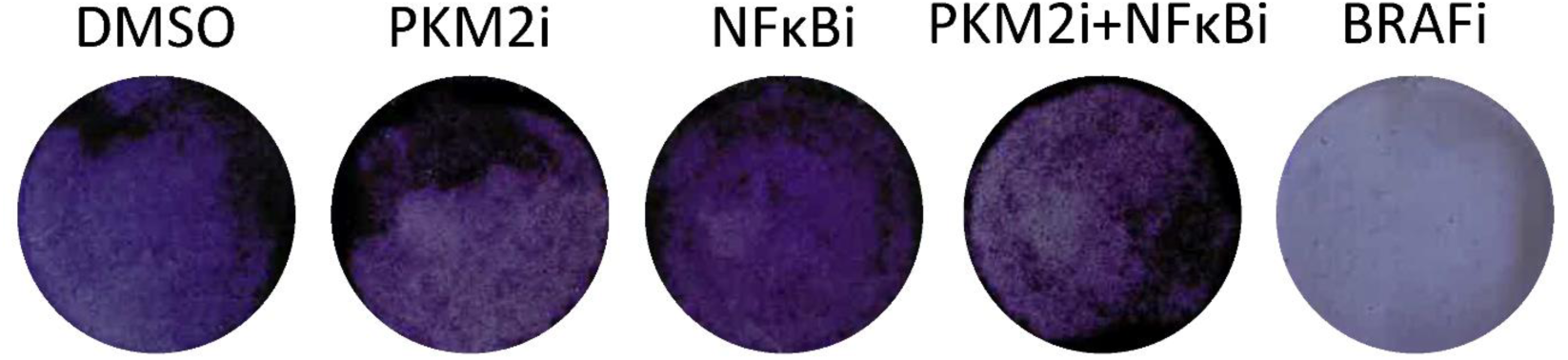
Short-term clongenic assay for 397 cells. M397 was treated with either DMSO control or PKM2i or NFKBi or PKM2i+NFKBi or BRAFi. No significant toxicity to the cells was observed for using PKM2i or NFKBi or combination of both.

